# Temporal chromatin and transcriptome dynamics driven by JUND and progesterone receptor binding in the pregnant mouse myometrium

**DOI:** 10.64898/2026.02.25.707818

**Authors:** Nawrah Khader, Andrew Duncan, Anna Dorogin, Oksana Shynlova, Jennifer A. Mitchell

## Abstract

The myometrium, the muscle layer of the uterus, undergoes profound phenotypic changes throughout pregnancy, labor, and postpartum, allowing for successful reproduction. Transcriptomic analysis of the murine myometrium revealed distinct gene expression signatures corresponding to these physiological stages, which reflect the dynamic remodeling processes occurring in the uterus during late gestation, term labor, and postpartum. These transcriptomic signatures at specific gestational points were accompanied by changes to the accessible chromatin landscape. Notably, increased chromatin accessibility was observed in the mouse myometrium at full term (gestational day 19) before the onset of active labor contractions. Accessible chromatin regions were bound by the transcription factor JUND, from the AP-1 family, and associated with progesterone receptor (PR) binding. Depletion of the progesterone receptor isoform B (PRB) from accessible chromatin at this prelabor stage implicates progesterone receptor isoform A (PRA) as a binding partner with JUND at prelabor and labor. During labor onset, accessible chromatin regions were associated with elevated production of enhancer RNAs and enriched in binding sites for transcription factors from the AP-1, SOX, and ETS families, implicating additional transcription factors in the labor process. Although PRB was strikingly absent from labor-associated accessible chromatin, it was found associated with histone H3 lysine 27 trimethylated (H3K27me3) repressed regions in late gestation and the postpartum period. These findings provide new insight into the dynamic transcriptional regulatory networks and chromatin-based mechanisms controlling gene expression in the myometrium across gestational stages providing new therapeutic targets for reproductive disorders.

## Introduction

The female uterus is a unique organ in the body capable of undergoing repeated cycles of massive growth, remodeling, and regression, a capacity that distinguishes it from any other adult tissue. During the process of pregnancy and term labor, the uterus undergoes structural and functional changes, in response to hormonal, mechanical and inflammatory stimuli (Lye *et al*., 2001; Khader *et al*., 2021). The functional muscle layer of the uterus (myometrium) consists mainly of smooth muscle cells which undergo profound phenotypic changes throughout gestation, from a quiescent state at early gestation, to a contractile state at the onset of labor. Over the duration of pregnancy, myometrial smooth muscle cells undergo hyperplasia and hypertrophy, allowing uterine growth and remodeling while maintaining a non-contractile state to successfully house the growing embryo(s) (Shynlova *et al*., 2009). In the mouse myometrium, high progesterone levels maintain quiescence of smooth muscle cells by repressing the expression of contraction-associated genes such as gap junction alpha 1 (*Gja1*, previously connexin 43), oxytocin receptor (*Oxtr*), prostaglandin receptor (*Pgfr*), prostaglandin endoperoxide synthase 2 (*Ptgs2*), and ion channels (Ou *et al*., 1997, 1998; Lye *et al*., 2001; Mitchell *et al*., 2004; Khader *et al*., 2021). By the end of gestation, the rapid decline of progesterone levels, coupled with the increase in mechanical tension on the uterine walls by the growing fetus(es), causes increased production of contraction-associated proteins (Shynlova *et al*., 2009) and a sterile inflammatory response (Shynlova *et al*., 2013a). These pathways induce the smooth muscle cell transition from a synthetic to a contractile phenotype causing the onset of labor. The completion of pregnancy, after successful labor, ends with postpartum uterine involution. Involution is characterized by the reduction of myometrial mass through apoptotic and autophagic pathways, the degradation of extracellular matrix (ECM) components, and the restoration of vascular and immune homeostasis (Nishinaka and Fukuda, 1991; Shynlova *et al*., 2004, 2009; Khader *et al*., 2021). Notably by day 4 postpartum (D4PP), the inflammatory response in the murine uterus subsides, ovarian cycles resume, with rising estrogen levels promoting endometrial regeneration and vascular remodeling (Hsu *et al*., 2014; Hackwell *et al*., 2023; Suarez *et al*., 2024).

Studies conducted in the human and mouse myometrium reveal that the transition from a non-laboring to a laboring state, involves profound changes in the transcriptome profiles of the myometrium (Shynlova *et al*., 2009; Mittal *et al*., 2010; Stanfield *et al*., 2019; Wu *et al*., 2020; Dotts *et al*., 2023). In mouse myometrium, gene transcription changes were observed between late gestation and labor (Shchuka *et al*., 2020), while in term non-laboring and laboring human myometrium increased transcription of many genes was found at labor onset (Khader *et al*., 2024). These changes in the transcriptome are thought to regulate the changing myometrial phenotypes during pregnancy, however, the molecular mechanisms that underpin these transcriptomic transitions remain poorly characterized.

Precise control of gene transcription is achieved by collections of non-coding regulatory regions and the transcription factors that bind to these regions (Khader *et al*., 2021; Tobias *et al*., 2021). Active regulatory regions in the genome are often more accessible than the surrounding chromatin, which allows transcription factors to bind and modulate gene expression. These regions, also known as “enhancers”, can contain many transcription factor binding motifs and recruit many transcription factors to communicate with target genes across great distances in the genome, regardless of their orientation (Liang *et al*., 2020; Singh *et al*., 2021). Active enhancers are often demarcated with modifications on neighboring histones such as monomethylation of lysine 4 in histone H3 (H3K4me1), as well as acetylation of lysine 27 in histone H3 (H3K27ac), and these features can be used to identify candidate enhancers (Heintzman *et al*., 2007; Creyghton *et al*., 2010; Rada-Iglesias *et al*., 2011; Moorthy *et al*., 2017; Tobias *et al*., 2021). These enhancers are also actively transcribed, producing short non-coding enhancer RNAs (eRNAs) (Kaikkonen *et al*., 2013; Mousavi *et al*., 2013; Bose *et al*., 2017; Tsai *et al*., 2018; reviewed in Sartorelli and Lauberth, 2020). Furthermore, enhancers in a “poised” state, contain both H3K4me1 (active) and repressive marks such as trimethylation of lysine 27 in histone H3 (H3K27me3) and therefore lack the mutually exclusive H3K27ac (Cao *et al*., 2002; Fischle *et al*., 2003). Collectively, these regulatory regions and their associated transcription factors form transcriptional regulatory networks capable of orchestrating complex cell behaviors; however, the transcriptional regulatory network that regulates transcriptional changes during the transition from myometrial smooth muscle cell quiescence to increased contractility, and the return to a non-contractile state remains largely unknown.

To understand how epigenetic regulation may contribute to these gene expression changes, Shchuka *et al*. (2020) conducted a genome-wide assessment of active histone marks, which demonstrated no changes in the active histone modifications, H3K27ac and H3K4me3, between pregnancy and labor in murine myometrium (Shchuka *et al*., 2020). This was in agreement with a later study conducted using human myometrium tissue from term non-laboring and laboring patients, which similarly found no major differences in active histone mark enrichment peaks for H3K27ac or H3K4me3 (Dotts *et al*., 2023). In this study, we explore an additional enhancer feature, chromatin accessibility, to identify changes at the chromatin level and present evidence for transcriptional regulators of phenotype-specific gene expression in murine myometrium during late gestation, term labor, and at day 4 postpartum, when the uterus has undergone sufficient repair to support a new pregnancy. Assay for transposase accessible chromatin paired with sequencing (ATAC-seq) of mouse myometrial tissue at different gestational time points demonstrated a global increase of chromatin accessibility prior to the onset of labor. Furthermore, differentially accessible regions display footprints of activator protein 1 (AP-1) factors at prelabor and active term labor, suggesting that the transition from relative quiescence to contractility requires a global opening of chromatin and AP-1 factor binding at these regions. This was corroborated through the observation of increased global enrichment of the AP-1 family member, JUND, at prelabor and labor, in comparison to earlier in pregnancy and the postpartum period, as revealed by chromatin immunoprecipitation paired with sequencing (ChIP-seq). The relatively fewer regions that are open earlier in pregnancy display a signature of progesterone receptor (PR) motifs, with elevated PRB isoform binding in the pregnant and postpartum tissues confirmed by PRB ChIP-seq. Progression to the postpartum state was concurrent with a global reduction in chromatin accessibility but did also reveal some novel regions that gain accessibility, and these are enriched in homeobox (HOX) sequence motifs. Coupled with increased expression of genes involved in wound healing processes, these data reveal the molecular changes involved in the restoration of the myometrium to a non-pregnant state during postpartum involution. Interestingly, differential ATAC-seq analysis revealed distinct gestational phenotype-specific clusters which are associated with increased gene expression suggesting that transcriptional changes are driven by changes in chromatin accessibility in the pregnant and postpartum myometrium. Finally, the assessment of transcribed intergenic regions, displaying accessibility at prelabor and labor, unveiled candidate labor enhancers which are associated with AP-1, SOX, and ETS transcription factor family motifs, which are likely important regulators of the contractility transcriptional regulatory network.

## Results

### Chromatin accessibility is dynamic in the mouse myometrium and associated with different transcriptional regulators across gestation

Previous studies have shown that the myometrial transcriptome changes profoundly between pregnancy and active labor. This change was not accompanied by changes to histone modifications associated with active chromatin, indicating that the myometrium is already at least partially epigenetically prepared for labor contractions several days before labor onset (Shchuka *et al*., 2020; Dotts *et al*., 2023). To examine whether the observed changes in gene transcript levels might be regulated by changes in chromatin accessibility, we employed ATAC-seq using mouse myometrium at late gestation, post-coitum day (D)15, D17, prelabor (PL; D19 not-in-labor), term labor (LAB; D19.5), and at two postpartum (PP) stages (D1PP/D20; D4PP/D23) (Fig 1A). Differences in chromatin accessibility were observed at transcription start sites (TSS), which globally exhibit increased accessibility at the prelabor stage, and dramatically reduced accessibility at day 1 postpartum (D1PP, Fig 1B). *Oxtr*, is a contraction-associated gene which plays a role in promoting contractility by responding to the hormone oxytocin (Kimura *et al*., 1996; Ou *et al*., 1998; Arrowsmith and Wray, 2014). We previously determined that *Oxtr* expression is increased during labor when compared to late pregnancy (D15) in the mouse myometrium (Shchuka *et al*., 2020). Here we observed an interesting trend in chromatin accessibility at the promoter of the *Oxtr* gene, and within the intron, with higher ATAC-seq signal intensity detected at prelabor and labor, compared to gestation (D15 and D17), and the postpartum period (D1PP and D4PP) (Fig 1C).

**Figure 1:**
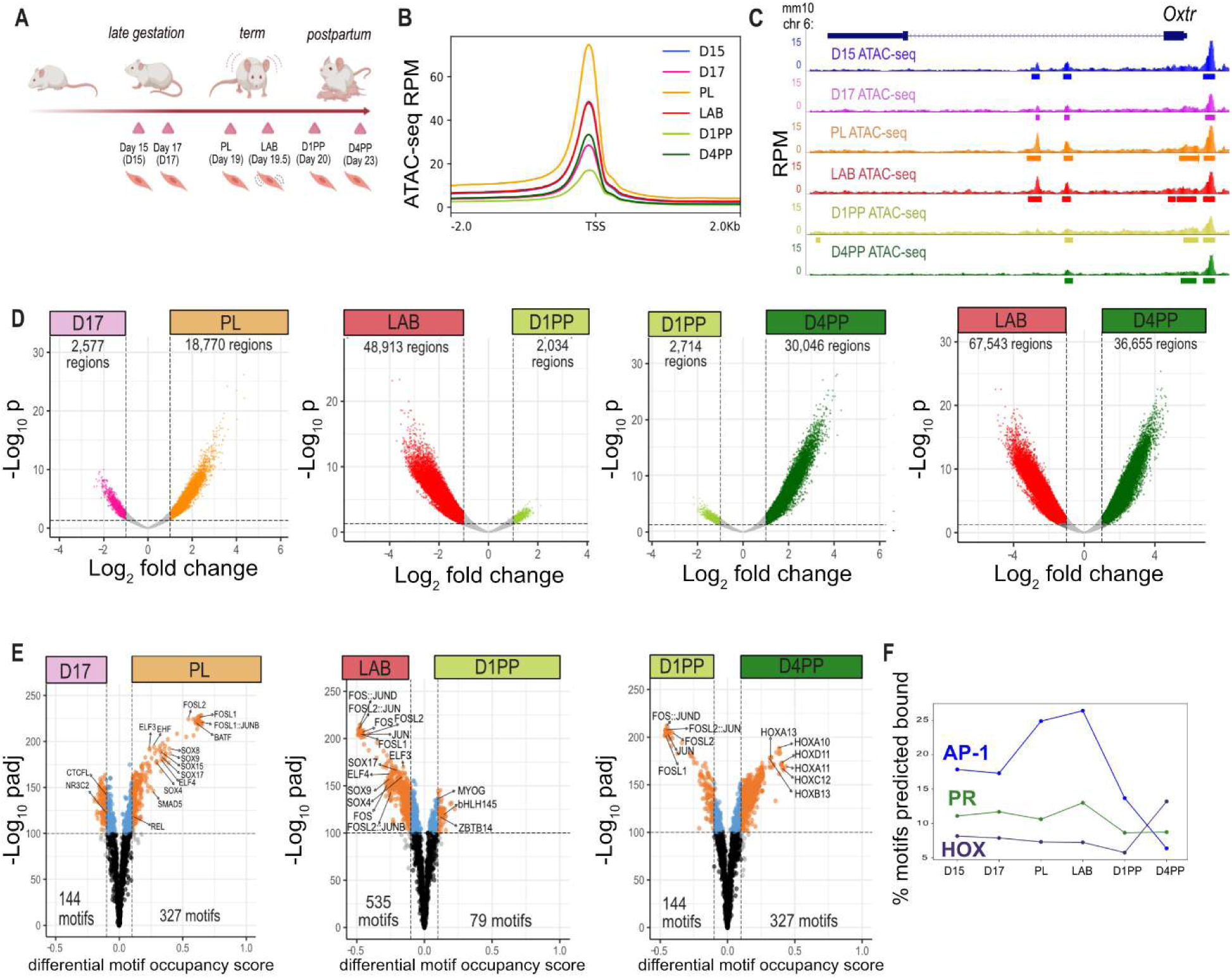
Changes in chromatin accessibility profiles and TFBS footprints throughout mouse gestation, labor, and postpartum. (A) Schematic showing the myometrium tissue samples used for ATAC-seq, obtained from mice at late gestation (D15 and D17), prelabor (D19; PL), term labor (D19.5; LAB) and postpartum (D1PP/D20, D4PP/D23). (B) ATAC-seq signal in reads per million (RPM) at gene promoters (TSS +/- 2kB). (C) Gene locus of labor-associated gene, *Oxtr*, displays chromatin accessibility signal in its gene promoter at all time points. (D) Pairwise comparisons in ATAC-seq reads. For all comparisons, |log_2_FC| ≥ 1 and FDR < 0.01 was used. (E) Pairwise comparisons of differentially enriched TFBS footprints. For all comparisons, differential motif occupancy score| ≥ 0.1 and FDR < 0.01 was used. (F) Temporal footprinting enrichment across gestational time points shown for representative AP-1 (FOSL1:JUND), progesterone receptor (PR; NR3C2) and HOX (HOXA11).

ATAC-seq revealed that accessible peaks are largely in intergenic (35-40%) and intronic regions (42-46%), consistent with a number of these peaks representing distal gene regulatory elements, with only 10-14% of peaks observed at gene promoters (Supplemental Fig 1A). Principal component analysis (PCA), as well as hierarchical clustering, revealed samples representing the same time point cluster together (biological replicates n ≥ 3), with the prelabor and labor samples clustering as one group (Supplemental Fig 1B/C). The pregnancy samples (D15/D17) were clustered in the PCA, although 2 of the D17 replicates were closer to the postpartum samples than the rest of the set. Pairwise differential chromatin accessibility analysis showed that there are no significant changes in chromatin accessibility between the two gestation time points (D15 and D17) or between prelabor and labor (Supplemental Fig 2A/B, Supplemental Tables 1 and 3) (**|**log_2_FC**|** ≥ 1, FDR < 0.01). In contrast to this, the transition from D17 to the prelabor state was accompanied by an increase in accessibility of close to 19,000 regions (Fig 1D, Supplemental Table 2). Comparison of labor to the postpartum samples (Fig 1E/F, Supplemental Table 4-6) indicated a closing of the labor-associated chromatin at D1PP, which was temporally separated from the gain in accessibility observed at D4PP, suggesting that a global closure of chromatin marks the successful completion of labor before the chromatin remodeling that returns the myometrium to the non-pregnant state. The most dramatic change in accessibility was seen by comparing labor and D4PP, with more than 67,000 regions that are accessible at labor losing accessibility by D4PP and over 36,000 regions gaining accessibility at D4PP (Fig 1G, Supplemental Table 6).

We next evaluated changes in the transcription factor binding site (TFBS) footprints within ATAC-seq peaks at these time points using TOBIAS (Bentsen *et al*., 2020). Footprints occur when transcription factors bind chromatin regions and prevent DNA cleavage by the transposase, resulting in a depleted ATAC-seq signal that can be used to infer transcription factor binding. Differential footprinting analysis using all open chromatin regions from each time point can identify changes at basepair resolution even in cases where there are no significant differences in accessibility at the level of whole peaks. Interestingly, despite no significant changes detected in chromatin accessibility between D15 and D17, differential footprinting analysis revealed 32 motifs enriched at D15 including CCCTC-binding factor (CTCF, **|**log_2_FC**|** ≥ 0.1, FDR < 0.01, Supplemental Table 7), a factor involved in chromatin architecture that acts to anchor interactions between gene promoters and distal regulatory elements (Ishihara *et al*., 2006; Phillips and Corces, 2009). In addition, 34 motifs were enriched at D17, including those bound by factors from the Sry-related HMG-box (SOX) family as well as progesterone receptor [PR; nuclear receptor subfamily 3 group C (NR3C)/androgen receptor (AR)] (**|**log_2_FC**|** ≥ 0.1, FDR < 0.01) (Supplemental Fig 2C, Supplemental Table 7).

When comparing D17 to prelabor, footprinting analysis revealed an enrichment of 144 motifs at D17, including PR (NR3C2/AR), CTCF and BHLH factors, and an enrichment of 327 motifs at prelabor which included factors belonging to the AP-1, SOX, and ETS families (Fig 1H, Supplemental Table 7). Despite the absence of significant changes in the locations of the genome that are accessible between the prelabor and labor time points, footprinting indicates that there are shifts in the most accessible basepair within these regions that suggest different transcription factors engage with the same regions in a dynamic way during this transition. Specifically, we identified an enrichment in 288 motifs in prelabor peaks, (predominantly AP-1, SOX, and HOX motifs), and 175 motifs enriched at labor (CTCF, NFKB, and NR3C/AR, or PR) (Supplemental Fig 2D). Differential footprinting comparison between labor and D1PP reveals that AP-1, SOX, and ETS family footprints are more highly enriched at labor, and depleted at D1PP indicating a loss of binding of these factors at D1PP as the labor-associated accessible chromatin becomes more closed (Fig 1I and Supplemental Fig 2E, respectively); however, when comparing D1PP and D4PP, AP-1 motifs were still enriched at D1PP compared to D4PP, indicating continual reduction in AP-1 motif accessibility during this time, as more clearly indicated by the time-course footprint analysis (Fig 1F). The most enriched motifs in D4PP accessible chromatin regions were members of the HOX family of transcription factors (Fig 1E, Supplemental Table 7), previously implicated in repressing the expression of specific contraction associated genes (Li *et al*., 2018).

Time-course footprinting reflects changes in the percentage of bound sites of select transcription factors throughout gestational time points (Fig 1F), which revealed motifs belonging to the AP-1 family (FOSL1:JUND shown as an example) exhibit the highest enrichment at labor compared to gestation (D15 and D17) as well as in comparison to postpartum (D1PP and D4PP). The prevalence of AP-1 motifs at accessible chromatin regions during labor onset is unsurprising as the regulatory role of AP-1 proteins in the laboring myometrium has been well characterized (Mitchell and Lye, 2002, 2005; Khanjani *et al*., 2012; Nadeem *et al*., 2018a; Peng *et al*., 2018; Shchuka *et al*., 2020; Khader *et al*., 2025). PR (NR3C2) motifs also showed an increase at labor compared to prelabor and postpartum, with a more subtle increase at D17 compared to D15 and prelabor (Fig 1F). In contrast, HOX motifs displayed increased accessibility predominantly at D4PP.

### RNA-seq reveals an increase in primary transcript levels occurs predominantly at prelabor

Having established that changes in chromatin accessibility accompany myometrial phenotypic transitions, we next examined gene expression at these gestational stages. Prior work assessing gene expression changes in the human myometrium looked at either non-pregnant vs term pregnant non-laboring (Wu *et al*., 2020), or term non-laboring vs laboring (Mittal *et al*., 2010; Chan *et al*., 2014; Stanfield *et al*., 2019; Dotts *et al*., 2023; Khader *et al*., 2025). Our earlier study compared gene expression in pregnant mouse myometrium (D15) to labor (Shchuka *et al*., 2020). Here we aim to determine transcriptional changes associated with the myometrium’s transition from late gestation to prelabor, labor, and then to postpartum. Since the ATAC-seq analysis suggests that the most significant changes happening at the chromatin level occur between D17 and prelabor as well as labor and D4PP, we conducted total RNA-seq, which allows for both exon and intron read analysis (Madsen *et al*., 2015), using murine myometrium tissues at D17, prelabor, labor, and D4PP (n = 3/group, Supplemental Fig 3A). Initial observation at the *Fos* gene locus identified elevated exon and intron reads at the prelabor and labor time points, compared to D17 and D4PP (Supplemental Fig 3B). Based on PCA and hierarchical clustering, conducted with the RNA-seq data, we observed clustering of the biological replicates from each time point, as expected (Supplemental Fig 3C and 3D). As in our prior transcriptional analysis of the mouse myometrium (Shchuka *et al*., 2020), and assessment of the transcriptomic profiles in the non-laboring and laboring human myometrium (Khader *et al*., 2025), which identified increased primary transcript levels associated with human term labor, we conducted differential pairwise gene expression analysis at both the exon and intron levels (Fig 2, Supplemental Tables 8 and 9, respectively). To understand, how the myometrium shifts from pregnancy to the prelabor stage, differential gene expression was conducted between D17 and prelabor which revealed an upregulation of 1,088 genes (Fig 2A, *left*), which included prominent labor associated- genes such as *Oxtr*, *Gja1*, matrix metalloproteinases 8 (*Mmp8*), 11 (*Mmp11*), and 28 (*Mmp28*), inflammatory proteins (Chemokine C-X-C motif ligands 1 [*Cxcl1*], 5 [*Cxcl5*], and 17 [*Cxcl17*]), as well as cell adhesion molecules (Carcinoembryonic antigen-related cell adhesion molecule 1 [*Ceacam1*], 5 [*Ceacam5*], 11 [*Ceacam11*]). Contrarily, 675 genes were found to be downregulated at prelabor compared to D17 which included genes involved in ECM architecture (Collagens type I alpha 2 [*Col1a2*], type XII alpha 1 [*Col7a1*], type XXVI alpha 1 [*Col26a1*]), and a number of water channels (Aquaporin 2 [*Aqp2*], 5 [*Aqp5*], 8 [*Aqp8*])(|log_2_FC| ≥ 1.5, *p* < 0.01). The set of genes found to be significantly upregulated or downregulated at prelabor compared to D17 (Supplemental Table 10) had approximately 22% overlap with genes previously found to be upregulated in murine myometrium during labor compared to D15 (Shchuka *et al*., 2020). Intron-based analysis at this time point reflected similar results to the exon-centric differential gene expression analysis with 1,214 genes upregulated at prelabor, and 659 genes found to be downregulated (Fig 2B, *left*). In fact, most genes (69%) upregulated at prelabor based on increased exonic reads, also displayed a significant increase in intron reads (Fig 2C, *left*, Supplemental Table 11), highlighting that transcriptional changes at labor-associated genes occurs at the prelabor stage.

**Figure 2:**
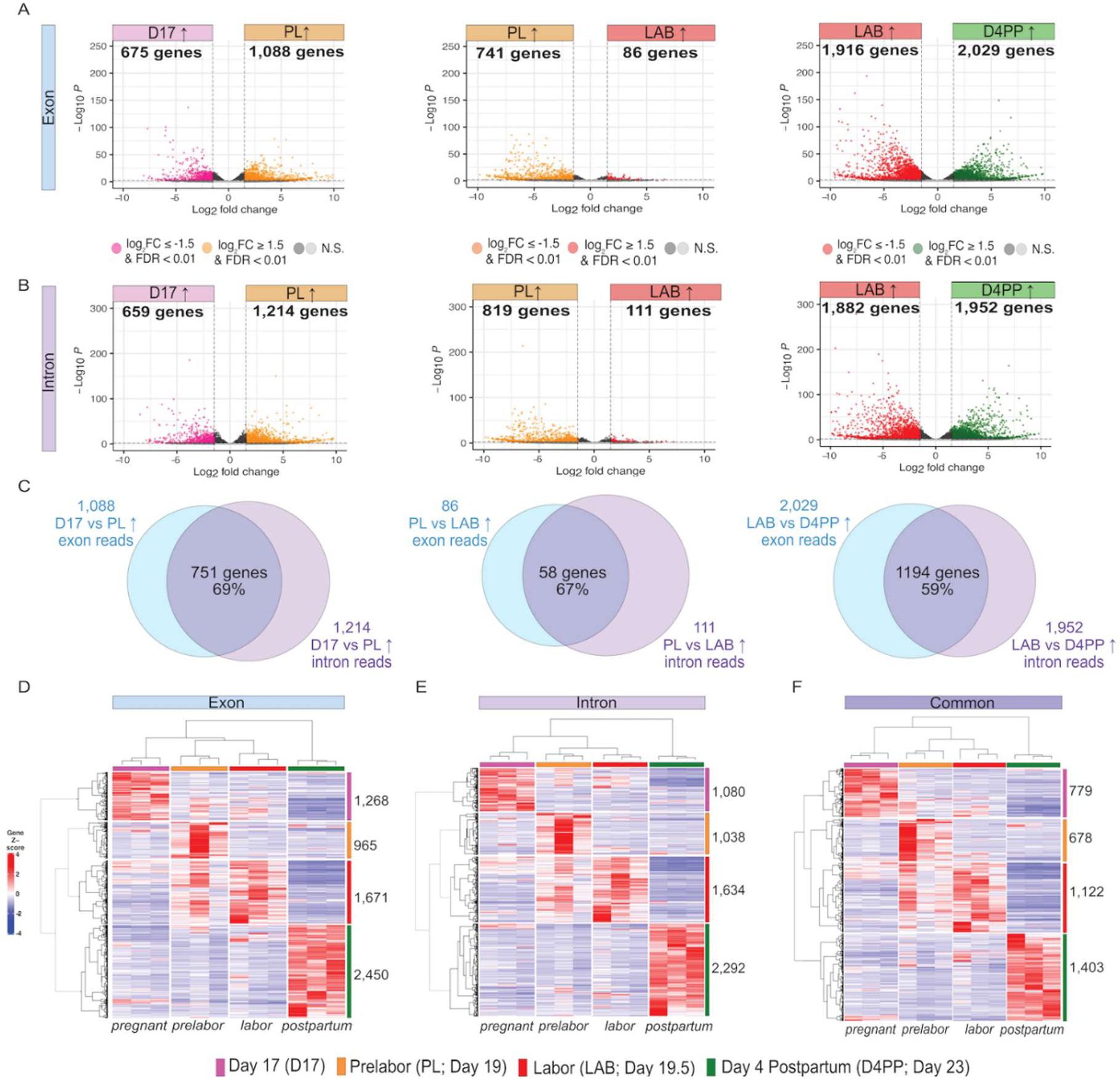
Primary transcript abundance changes accompany the transitions between gestational stages in the myometrium. RNA-seq volcano plots highlighting differential gene expression between D17 and PL at the exon (A) and intron (B) level, with Venn diagram displaying the overlap between the number of genes differentially expressed at the exon and intron level (C). Comparisons were done between D17 and PL, PL and LAB, and between LAB and D4PP. For all comparisons, |log_2_FC| ≥ 1.5, FDR < 0.01 was used. Heatmap demonstrating hierarchical clustering of gene groups based on RNA expression changes between pregnant (D17), prelabor (PL), labor (LAB), and postpartum (D4PP) at the exon (D), intron (E). Genes that were differentially expressed at both the exon and intron level were termed as “merged” and used to perform hierarchical clustering of gene groups (F).

**Figure 3:**
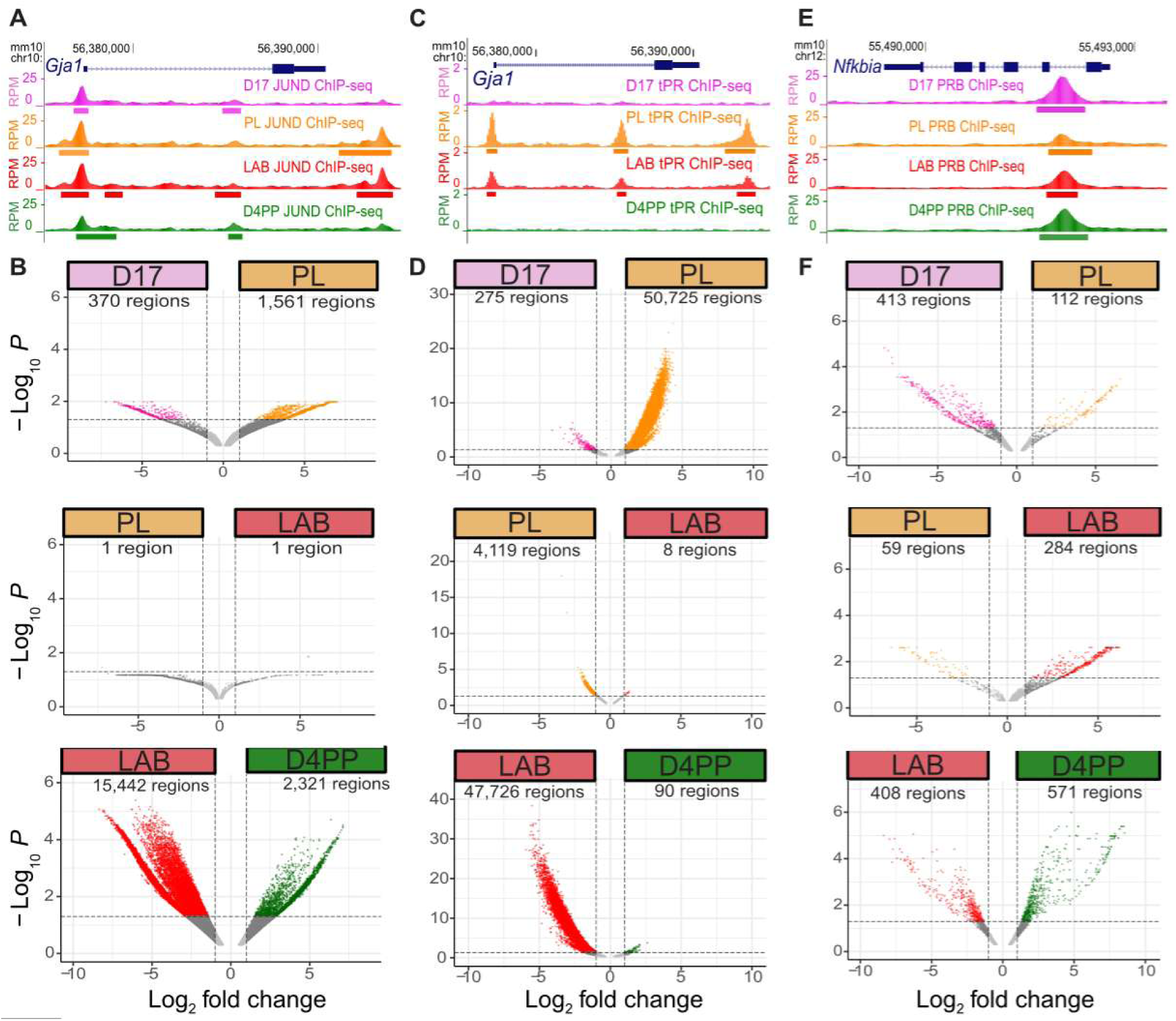
JUND and PR isoforms display dynamic binding throughout gestation in the myometrium. (A) JUND ChIP-seq (reads per million; RPM) at the *Gja1* gene locus during late gestation (D17), prelabor (PL), labor (LAB), and postpartum (D4PP). (B) Volcano plots of pairwise comparisons depicting differential JUND enrichment between D17 and PL, PL and LAB, and LAB and D4PP. For all comparisons, |log_2_FC| ≥ 1 and FDR < 0.05 was used. (C) Total PR (tPR) ChIP-seq at the *Gja1* gene locus. (D) Volcano plots of pairwise comparisons depicting differential tPR enrichment between D17 and PL, PL and LAB, and LAB and D4PP. For all comparisons, |log_2_FC| ≥ 1 and FDR < 0.05 was used. (E) PRB ChIP-seq at the *Nfkbia* gene locus. (F) Volcano plots of pairwise comparisons depicting differential PRB enrichment between D17 and PL, PL and LAB, and LAB and D4PP. For all comparisons, |log_2_FC| ≥ 1 and FDR < 0.05 was used.

Despite exhibiting minimal differences in the accessible chromatin landscape, transcriptomic analysis revealed that the transition from prelabor to labor is accompanied predominantly by a decrease in gene expression, with 741 genes that are downregulated and 86 genes that are upregulated at labor, in comparison to prelabor (Fig 2A, *middle*). These results were corroborated by intron RNA-seq analysis that revealed a downregulation of 819 genes and an upregulation of 111 genes from prelabor to labor (Fig 2B, *middle*), with 67%, or 58 upregulated genes, overlapping the exon read analysis (Fig 2C, *middle*). Amongst the genes exhibiting increased expression at prelabor is Myosin XVB (*Myo15b*), which is involved in the acto-myosin complex and predicted to enable cytoskeletal motor activity. Genes upregulated at labor include *Ptgs2*, *Cxcl2*, and *Mmp11,* all previously shown to have higher expression in the laboring murine myometrium compared to late gestation (Shchuka *et al*., 2020). These data shed light on the dynamic changes in myometrial transcription during the transition from a relatively quiescent state, at prelabor, to a contractile state, at labor onset.

Finally, similar to the ATAC-seq data, the largest change in transcriptomic profiles was observed between labor and D4PP samples with exon-based analysis revealing a significant upregulation of 2,028 genes, and downregulation of 1,916 genes at D4PP vs labor (Fig 2A, right). This trend was mirrored in the intron-based differential gene expression analysis which revealed an upregulation of 1,952 genes, and a downregulation of 1,882 genes in the D4PP vs labor (Fig 2A, *right*). Genes exhibiting higher expression at labor compared to D4PP included those previously observed in the laboring murine myometrium such as *Gja1, Fos,* E74 like ETS transcription factor 3 (*Elf3*), *Ceacam1*, and Sry-related box transcription factor 7 (*Sox7*), and 9 (*Sox9*) (Shchuka et al., 2020). Upregulated genes in the postpartum (D4PP) myometrium included genes involved in ECM architecture (i.e. Collagens type XXII alpha 1 [*Col22a1*], type XXVI alpha 1 [*Col26a1*]), and transcription factor encoding genes of the HOX family (i.e. homeobox protein A9 [*Hoxa9*], A10 [*Hoxa10*], A11 [*Hoxa11*], D9 [*Hoxd9*], D11 [*Hoxd11*]), and those previously more associated with D15 murine myometrium (transcription factor 23 [*Tcf23*] and Zinc Finger And BTB Domain-Containing Protein 16 [*Zbtb16*]) (Shchuka *et al*., 2020). Overlapping gene set analysis confirmed that, similar to the previous pairwise comparisons, the majority of genes exhibiting significant exon read accumulation (59%) also contain increased intron reads (Fig 2C, *right*), underscoring the gestational stage-specific changes in primary transcript levels in the laboring and postpartum myometrium. Hierarchical clustering uncovered separate subsets of genes that cluster based on differential gene expression comparisons at the exon (Fig 2D, Supplemental Table 12) and intron level (Fig 2E, Supplemental Table 13) as well as a comparison of genes, termed “common”, that are differential at both, the exon and intron level (Fig 2F, Supplemental Table 14). These results demonstrate that the phenotypic transition of the myometrium between different gestational stages is due in large part to changes in primary transcript levels. These collective findings strongly suggest that substantial transcriptional activity occurs as the myometrium transitions from late gestation (D17) to the prelabor stage in preparation for labor onset, from prelabor to the contractile phase of labor, and finally at the end of the labor state during the transition to the postpartum involution and remodeling period (D4PP). These transcriptional changes are likely driven by the action of the different transcription factor families identified by the ATAC-seq footprinting analysis at each of these stages (Fig 1).

Gene ontology analysis reveals that the genes expressed at each stage are involved in distinct processes. Specifically, the genes expressed at D17 are involved in processes including ion transport which was previously observed in genes expressed in the pregnant mouse myometrium at D15 (Supplemental Fig 4A) (Shchuka *et al*., 2020). Genes demonstrating increased expression prelabor were involved in muscle contraction, while those at labor were involved in inflammatory processes (Supplemental Fig 4B and 4C, respectively). This supports previous suggestions that genes required in preparing the myometrium for contractions are expressed before labor onset and maintained until labor begins, whereas sterile inflammatory processes in the myometrium happen acutely during labor to induce contractions (Bollopragada *et al*., 2009; Dotts *et al*., 2023). Finally, the genes with elevated expression postpartum are involved in negative regulation of cell adhesion, peptidase activity, and wound healing (Supplemental Fig 4D). These alterations are indicative of shutdown of the laboring process and the completion of postpartum uterine involution, which requires MMPs to degrade the ECM for tissue reorganization as the myometrium returns to a non-pregnant state (Shynlova *et al*., 2009).

**Figure 4:**
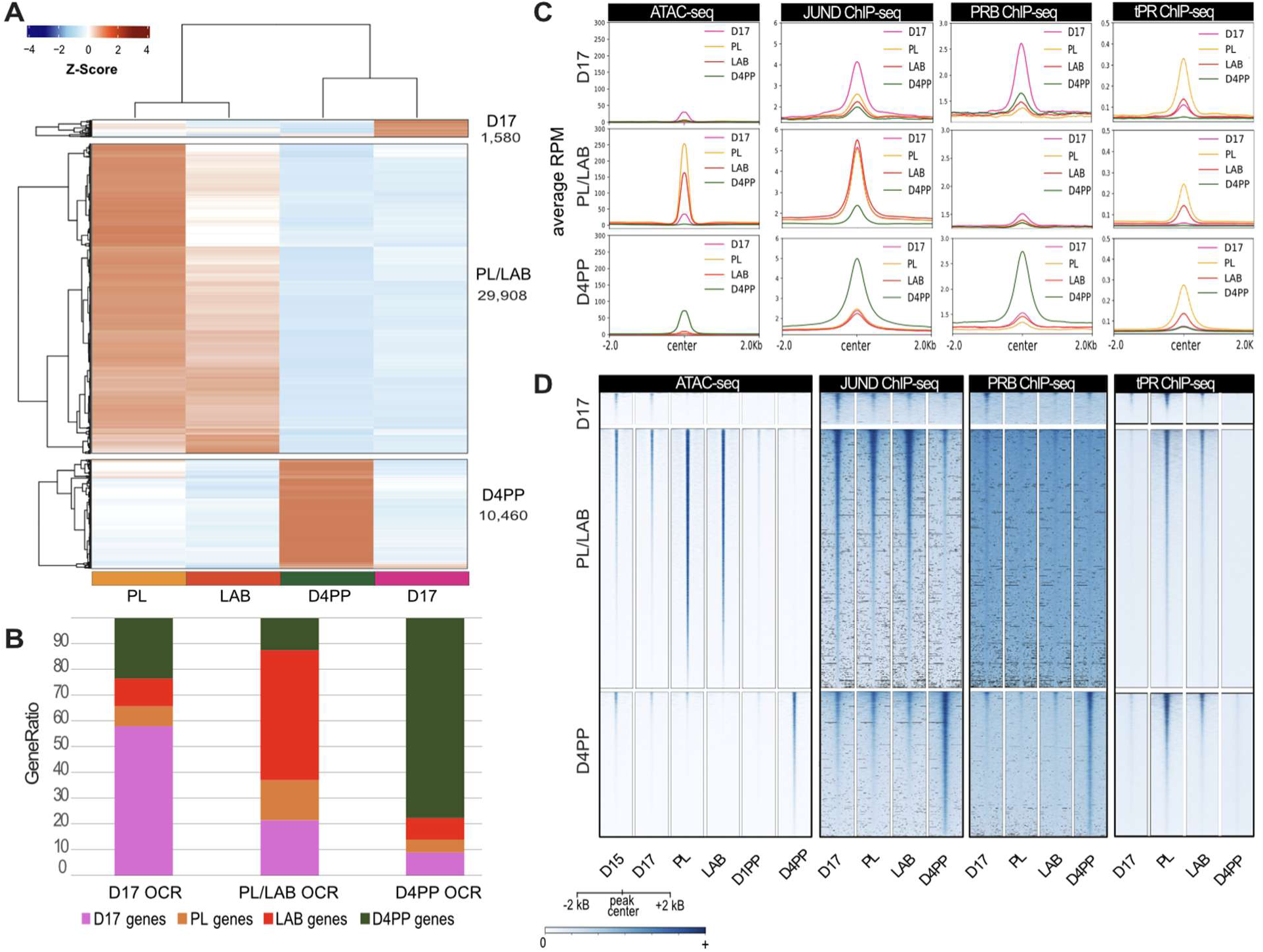
Myometrial phenotype-specific open chromatin regions drive expression of genes required at each gestational stage. A) Heatmap depicting differential open chromatin region clusters at late gestation (D17), term (prelabor; PL and labor; LAB), and postpartum (D4PP). B) Stacked bar graph depicting the distribution of genes that are associated with promoter-specific peaks in open chromatin region (OCR) clusters at each gestational stage, including pregnancy (D17), term (PL/LAB), and postpartum (D4PP). C) Profile plots displaying ATAC-seq signal, JUND ChIP-seq signal, PRB and total PR (tPR) ChIP-seq signal at each gestational stage ATAC-seq cluster. D) Heatmap plots displaying ATAC-seq signal, JUND ChIP-seq signal, PRB and total PR (tPR) ChIP-seq signal at each gestational stage ATAC-seq cluster.

### JUND and PR exhibit dynamic binding throughout gestation in the myometrium

Since AP-1 factors have long been implicated as regulators of labor initiation, and ATAC-seq data revealed an over-representation of AP-1 motifs at the prelabor and labor stages, we next investigated the genome regions bound by the AP-1 family member, JUND. Prior work, including genome-wide analysis from the human myometrium, revealed JUND as a factor enriched at term, that may regulate the contraction-associated gene *Gja1* (Mitchell and Lye, 2005; Nadeem *et al*., 2016; Khader *et al*., 2025). For this reason, we conducted ChIP-seq for JUND using mouse myometrium tissues from D17, prelabor, labor, and D4PP (n=2 /time point). Motif enrichment of JUND ChIP-seq peaks confirmed an over-representation of AP-1 motifs at all time points (Supplemental Tables 15-18). PCA and hierarchical clustering confirms correlation between replicates belonging to the same time point (Supplemental Fig 5A and 5B). Initial assessment of *Gja1* demonstrates the gene promoter is bound by JUND at all time points, with additional peaks found downstream of the gene at prelabor and labor (Fig 3A).

**Figure 5:**
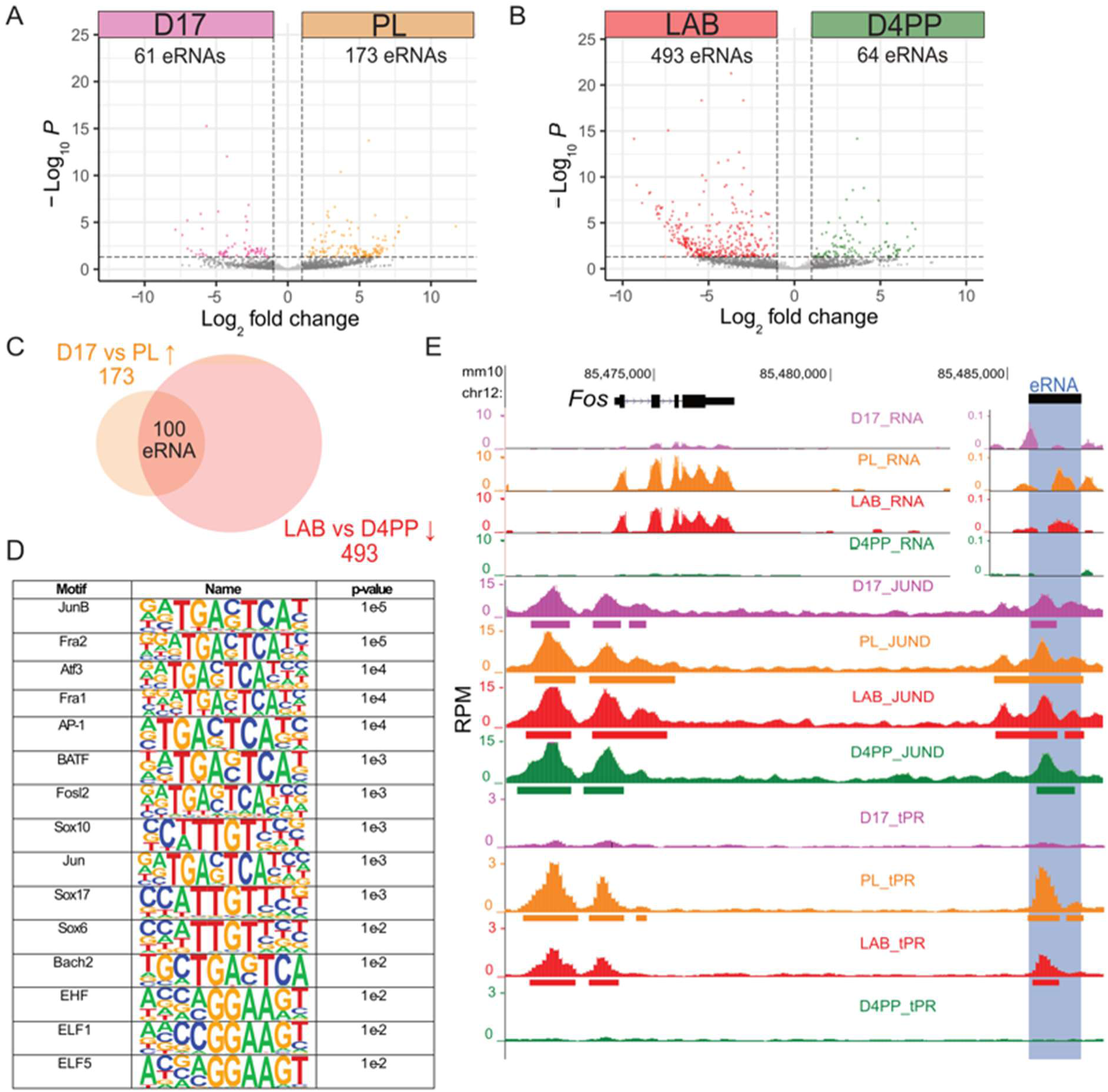
Increased enhancer RNAs (eRNAs) at prelabor and labor are associated with AP-1, SOX, and ETS factors. Volcano plot displaying differential eRNA expression between A) day 17 (D17) and PL (prelabor); B) between LAB (labor) and D4PP (day 4 postpartum) at intergenic differential open chromatin regions from the PL/LAB cluster. For all comparisons, |log_2_FC| ≥ 1, FDR < 0.05 was used. C) Venn diagram depicting overlap of eRNAs gained at PL in comparison to D17 with eRNAs gained at LAB in comparison to D4PP. D) Motif analysis of overlapping regions displaying increased differential eRNA expression at PL and LAB. E) JUND and PL/LAB PR bound region downstream of labor-associated gene, *Fos*, displays increased eRNA at PL and LAB.

Similar JUND enrichment patterns were observed at genomic features across time points, with the largest number of JUND peaks found at intronic and intergenic regions (40-45% for each), consistent with the role of JUND as a transcription factor acting predominantly at distal cis-regulatory regions (Supplemental Fig 5C). Pairwise comparisons between time points confirm increased JUND accumulation at prelabor as compared to D17 (1,561 regions gained, Fig 3B, Supplemental Table 19) (|log_2_FC| ≥ 1, FDR < 0.05). There were no significant changes in JUND binding between prelabor and labor (Fig 3B, Supplemental Table 20). However, the largest shift in JUND enrichment occurs between labor and D4PP, with 15,442 labor peaks lost and 2,321 peaks gained at D4PP (Fig 3B, Supplemental Table 21). This differential enrichment analysis highlights a role for JUND as a transcriptional regulator in preparing murine myometrial cells for the onset of labor, similar to human myometrium (Khader *et al*., 2025), and additionally highlights the loss of JUND engagement in the postpartum remodeling phase as the myometrium returns to a non-pregnant state.

Through seminal work done in the *Gja1* locus, and prior findings in the human myometrium, PR isoforms have been implicated to interact with AP-1 complexes in regulating the expression of *Gja1* with the PRB isoform acting as a transcriptional repressor whereas PRA acts as an activator (Nadeem *et al*., 2016; Khader *et al*., 2024). To broaden our understanding of the role for PR isoforms as the myometrium transitions between different phenotypic states, we conducted ChIP-seq for total PR (tPR, antibody that recognizes both PRA and PRB) and PRB using mouse myometrium tissues from D17, prelabor, labor, and D4PP (n = 2/time points). Antibodies are not available for the PRA isoform as the PRA sequence is entirely contained within the larger PRB isoform. PRB and tPR PCA and hierarchical clustering revealed correlation between replicates belonging to the same time points (Supplemental Fig 6). Assessment of tPR peaks at genomic features revealed an increase in the number of peaks at gene promoters in the prelabor and labor samples (13-14%) compared to D17 and D4PP (4-5%) with a coordinate decrease in the proportion of intergenic peaks (46% at D17 to 24-27% at PL/LAB, Supplemental Fig 6F). PRB peak locations were more consistent across gestation with the largest enrichment found at intronic and intergenic regions (65-71% intergenic, 25-21% intronic) (Supplemental Fig 6C).

Assessment of tPR at the *Gja1* locus demonstrates the gene promoter, intron and the downstream region, each corresponding to JUND bound regions (Fig 3A), are also bound by PR only in the prelabor and labor time points (Fig 3C) suggesting this is primarily PRA binding, as PRA expression is higher than PRB in the labor state (Mesiano *et al*., 2002; Merlino *et al*., 2007). In fact, PRB ChIP-seq revealed no peaks overlapping *Gja1* at any of the time points investigated (Supplemental Fig 6G). In contrast to this, examination of the nuclear factor-kappa-B inhibitor alpha gene (*Nfkbia)* locus, involved in mediating the inflammatory response at labor (Chan *et al*., 2014), displays PRB binding in the first intron at all time points (Fig 3E), with increased signal intensity at the D17 and postpartum stages when PRB expression is higher than PRA (Mesiano *et al*., 2002; Merlino *et al*., 2007).

Both tPR and PRB ChIP-seq revealed dynamic binding across the gestational time points (|log_2_FC| ≥ 1, FDR < 0.05), with some interesting differences (Fig 3D/F). Total PR binding revealed a large gain occurs at prelabor (50,725 regions) with a subsequent loss at D4PP (47,726 regions; Fig 3D, Supplemental Table 22-24), corresponding to expression dynamics of PRA (Mesiano *et al*., 2002; Merlino *et al*., 2007). In contrast to the observations made for tPR, PRB displayed fewer regions bound genome wide and fewer regions with dynamic binding (Fig 3F, Supplemental Table 25-27). Similar to JUND, the largest shift in PRB dynamics is exhibited between the labor and D4PP myometrium, with a loss of PRB binding at 408 regions and a gain in PRB binding at 571 regions (Fig 3F, Supplemental Table 27).

Motif enrichment of tPR ChIP-seq peaks revealed that at D17 and D4PP the most enriched motifs are nuclear receptors including for PR, whereas at prelabor and labor the most enriched motifs are those bound by the AP-1 family with PR displaying less prominent enrichment (Supplemental Tables 28-31), which aligns with our prior motif assessment of PR occupied regions in the human myometrium (Khader *et al*., 2024). Motif enrichment for PRB ChIP-seq peaks revealed motifs for nuclear receptors including for PR are enriched at all time points (Supplemental Tables 32-35). Additionally, motif analysis for PRB at prelabor and labor demonstrated an additional enrichment of AP-1 factor motifs. Interestingly, PRB bound regions at D4PP also displayed an overrepresentation of the HOX motifs observed in our assessment of open chromatin regions obtained at this time point (Fig 1F). This is consistent with PR binding chromatin through interactions with other factors, which has been observed in other cell types (Rubel *et al*., 2012).

### Regions with differential chromatin accessibility are associated with myometrial phenotype-specific gene expression and dynamic JUND /PR engagement

In order to better understand the dynamics of transcription factor binding at regions with differential chromatin accessibility, we performed an integrative analysis using ATAC-seq and transcription factor ChIP-seq data. Since differential ATAC-seq assessment showed the largest variation in chromatin accessibility between D17 and prelabor, or labor and D4PP, we used the differentially accessible regions obtained from pairwise comparisons (Fig 1) to further scrutinize regions that display time point-specific chromatin accessibility (Fig 4A, Supplemental Table 36). Through this analysis, we observed 1,580 open chromatin regions specific to D17; 29,908 open chromatin regions at prelabor and labor (noted as PL/LAB); and 10,460 open chromatin regions at D4PP.

Assessment of gene promoters at these time points revealed that open chromatin regions at D17 are enriched in promoters of genes with higher expression at D17, whereas PL/LAB open chromatin regions are enriched in promoters of genes with higher expression at labor, and D4PP open chromatin regions are enriched in promoters of genes with higher expression at D4PP (Fig 4B, Supplemental Table 37, exon/intron gene list from Fig 2F/ Supplemental Table 14). In addition, a similar trend is observed when the intronic and intergenic peaks are assigned to the closest gene, with PL/LAB accessible chromatin displaying an enrichment of genes expressed at increased levels in labor (Supplemental Fig 7). This analysis suggests that changes in chromatin accessibility are major regulators of the time point-specific transcriptional surges in the myometrium.

To assess changes in transcription factor binding at time point-specific accessible chromatin regions, we conducted an enrichment analysis of JUND, tPR, and PRB at distal differential accessible chromatin region clusters displayed as profile plots and heatmaps centered on the ATAC-seq peaks (+/-2 kb, peaks that overlap a gene TSS +/- 2kb were removed) (Fig 4C and 4D, respectively). Through this analysis at D17 and D4PP accessible chromatin clusters, we detected binding of JUND and PRB. However, at the PL/LAB cluster regions we observed the binding of JUND, and strong tPR signal in the absence of PRB binding, pointing to PRA as predominantly engaging these regions. Interestingly, whereas the PL/LAB open chromatin regions display the largest ATAC-seq enrichment at PL/LAB stage, they are accessible at late gestation (D15 and D17) and bound by JUND at these time points (Fig 4C), indicating JUND pre-marks the regions that gain accessibility and PRA binding at the PL/LAB stage.

### Labor-associated enhancers demonstrate enrichment of AP-1, SOX, and ETS motifs

Our observation that at D17, JUND pre-binds regions of the genome that become more accessible at the prelabor stage and display increased binding in the tPR data, likely reflecting PRA association, suggests that additional factors bind these regions to activate labor-specific gene transcription. Given that a substantial portion (∼40%) of these peaks lie within intergenic regions, we were interested in dissecting their regulatory function by analysis of enhancer RNA (eRNA) at these gestational stages. Filtering the D17- and D4PP-specific intergenic open chromatin regions resulted in a small number of peaks with very few eRNAs detected. Additionally, as we were primarily interested in identifying enhancer regions that are drivers of the laboring process in the myometrium, we focused on the 10,613 intergenic regions in the PL/LAB specific open chromatin region cluster. Differential eRNA expression analysis determined that 61 eRNAs are downregulated and 173 eRNAs are upregulated between D17 and prelabor (Fig 5A, Supplemental Table 38); 18 eRNAs are downregulated and 4 eRNAs are upregulated between prelabor and labor (Supplemental Fig 7, Supplemental Table 39); and 493 eRNAs are downregulated and 64 eRNAs are upregulated between labor and D4PP (Fig 5B, Supplemental Table 40) (|log_2_FC| ≥ 1, FDR < 0.05). This analysis suggested that the greatest change to enhancer activity occurs as the myometrium transitions from pregnant D17 to prelabor, and then from labor to D4PP. To further analyze the enhancer regions in the mouse myometrium that are associated with labor, regions displaying elevated eRNAs at prelabor compared D17 and at labor compared to D4PP were overlapped which revealed 100 potential enhancer regions (Fig 5C, Supplemental Table 41). Motif enrichment analysis of these regions identified an over-representation of transcription factors belonging to the AP-1, SOX, and ETS families (Fig 5D, Supplemental Table 42).

One of these candidate regions downstream of the *Fos* gene locus demonstrates significantly increased eRNA expression at prelabor and labor in comparison to D17 and D4PP, respectively (Fig 5E). This region is bound by JUND and displays peaks of total PR, but not PRB, at prelabor and labor, as well as RNAPII, H3K27ac, and H3K4me3 signatures (Shchuka *et al*., 2020). Moreover, comparison of this region to the human genome (hg38) uncovers a putative labor-specific enhancer that displays elevated eRNA, H3K27ac, and is bound by JUND in the human myometrium (Khader *et al*., 2024). Interestingly, this region was previously highlighted in rat neuronal cells as an enhancer that drives the expression of *Fos* (Carullo *et al*., 2020). Conservation of this region across three different species, chromatin accessibility, RNAPII recruitment, H3K27ac, JUND, and PR transcription factor binding are all classical hallmarks of enhancers, implicating this region, and specifically JUND and PR, in regulating transcription of the *Fos* gene during active labor.

### H3K27me3 is elevated in the prelaboring myometrium

Although, chromatin accessibility profiles between prelabor and labor exhibited no significant differences, transition from a prelaboring to laboring stage is marked by transcriptional changes (Fig 2). For this reason, we questioned whether there are any changes in the presence of repressive histone marks, such as trimethylation of histone H3 on lysine 27 (H3K27me3) at regulatory regions, that could explain maintained repression in the myometrium before labor onset. We conducted H3K27me3 ChIP-seq at D17, prelabor, labor and D4PP. Inspection of the *Myo15b* gene, a prelabor upregulated gene involved in contractility in the myometrium, revealed H3K27me3 enrichment at the promoter at all time points, except for prelabor (Fig 6A). Hierarchical clustering and PCA confirm that replicates belonging to the same time point cluster together, and there is a distinct separation of the samples collected from different time points (Supplemental Fig 8B/C). Peak annotation revealed that the majority of H3K27me3 signal is enriched at intergenic and intronic regions (65-71% intergenic, 26-32% intronic) with only ∼1% in promoter and exon regions (Supplemental Fig 8D).

**Figure 6:**
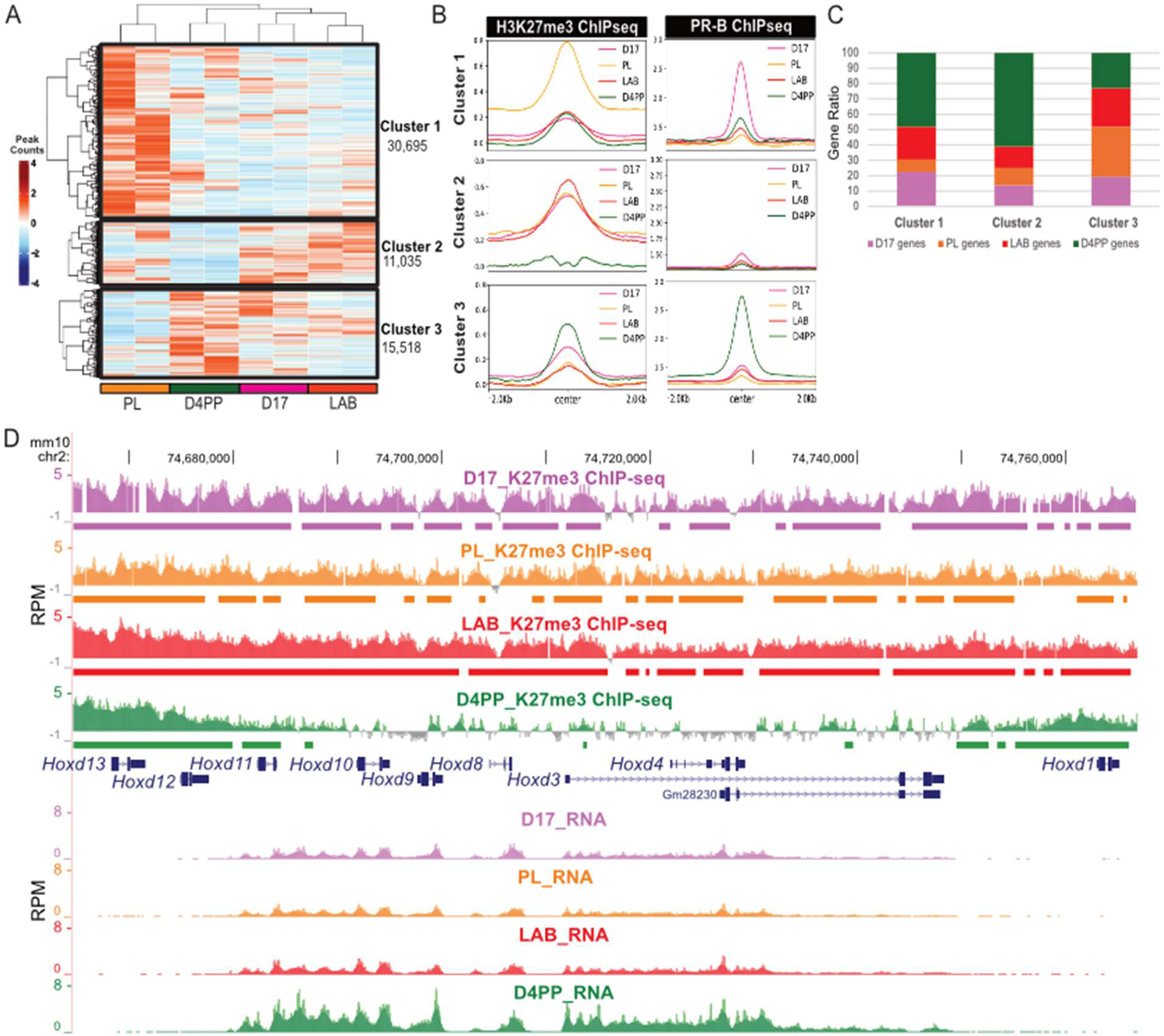
H3K27me3 modified regions in pregnant myometrium are associated with gene expression postpartum. A) Heatmap displays differential H3K27me3 peaks that forms three clusters. B) Profile plots exhibiting the H3K27me3 signal, ATAC-seq signal, JUND ChIP-seq signal, and PRB ChIP-seq signal at the center of peaks (+/- 2kB) from each cluster. C) Stacked bar graph depicting the distribution of time point-specific genes that are associated with promoter peaks in each cluster. Genes demonstrating time point specific expression common at both the intron and the exon level were used (from Fig 2J,F). D) H3K27me3 ChIP-seq and RNA-seq reads (reads per million; RPM) at the *Hoxd* locus demonstrates H3K27me3 erosion postpartum (D4PP) over the postpartum upregulated *Hoxd* genes.

Differential analysis was conducted among the time point comparisons as done for RNA-seq and transcription factor ChIP-seq (Supplemental Fig 8E-G, Supplemental Tables 43-45). To investigate myometrial H3K27me3 dynamics, we conducted a clustering analysis of these differential regions (Fig 6A). Hierarchical clustering, based on peaks with differential enrichment (|log_2_FC| ≥ 1, FDR < 0.01, Supplemental Table 46) revealed a subset of 30,695 peaks (Cluster 1; prelabor repressed) that show elevated H3K27me3 signal at prelabor only. Furthermore, a second set of 11,035 peaks (Cluster 2; pregnancy repressed) displays H3K27me3 enrichment at all stages except for D4PP. The final set of 15,518 peaks (Cluster 3; postpartum repressed) exhibits a higher H3K27me3 signal in the D4PP samples compared to the other gestational time points, but most notably the prelabor stage. This differential analysis suggests that the largest gain in the repressive H3K27me3 marks occurs at prelabor compared to the other gestational time points.

Next, we examined the overlap in feature enrichment for H3K27me3, ATAC-seq, JUND, tPR and PRB signals at these three clusters (Fig 6B and Supplemental Fig 8H). Surprisingly, we saw virtually no overlap between these dynamic H3K27me3 regions and accessible chromatin or JUND and tPR binding at any time point (Supplemental Fig 8H), indicating that H327me3 is not contributing to the repression of enhancers prior to later activation, as has been observed for developmental transitions (Zenk *et al*., 2017). Interestingly, however, the prelabor repressed cluster demonstrates elevated H3K27me3 enrichment at prelabor as well as increased PRB enrichment at D17, indicating PRB pre-marks regions for later repression. This PRB enrichment was also observed in the D4PP repressed cluster, where H3K27me3 and PRB enrichment were both observed at D4PP (Fig 6C). Given that PRB, has been shown to interact with transcriptional co-repressors, it is unsurprising to see PRB enrichment at regions associated with the repressive H3K27me3 modification (Dong *et al*., 2009; Nadeem *et al*., 2016), however, this pre-marking by PRB of regions that later become repressed at prelabor has not been reported previously.

Interestingly, the gene promoters associated with these clusters reveal distinct expression patterns (Fig 6C, Supplemental Table 47). The prelabor and pregnancy repressed clusters 1 and 2, which exhibit the lowest H3K27me3 enrichment at D4PP, are mostly linked with D4PP expressed genes [51% out of 452 genes in total (cluster 1) or 59% out of 487 genes in total (cluster 2)]. The D4PP repressed cluster 3, displaying higher H3K27me3 signal at D4PP, is associated with fewer D4PP genes but a higher percentage of prelabor-related genes (32%) in comparison to the other two clusters (6.9% and 11.3%, respectively). These findings cumulatively highlight that while H3K27me3 does not regulate the observed changes in chromatin accessibility or JUND binding throughout gestation in the myometrium, it appears to have an important role during pregnancy by repressing postpartum-expressed genes, and the repression of labor-associated genes during postpartum involution in cooperation with PRB. A striking example of this repression of postpartum-expressed genes can be seen at the *Hoxd* locus which is covered by H3K27me3 in all pregnancy time points (Fig 6D). In the postpartum myometrium we observed a striking erosion of the H3K27me3 signal over part of this locus. Notably, the postpartum erosion domain corresponds to the genes in this locus with increased expression postpartum compared to the pregnancy time points (*Hoxd11*, *10*, *9*, *8*, *3*, *4*), whereas genes that remain covered by H3K27me3 (*Hoxd13*, *12*, *1*) are not expressed (Fig 6D). This finding links a chromatin regulatory mechanism to the increased expression of *Hoxd* genes explaining the increase in HOX motif accessibility observed in the postpartum myometrium by ATAC-seq (Fig 1).

## Discussion

Our findings suggest that chromatin accessibility and transcriptomic changes in mouse myometrium from late pregnancy to the postpartum period are driven by the coordinated action of different transcription factors. In particular, AP-1 transcription factor motifs dominate in accessible chromatin regions at prelabor and labor, implicating AP-1 factors in inducing contraction associated gene expression for labor onset, whereas HOX motifs appear to be involved in new accessible chromatin postpartum. JUND binding occurs at the prelabor/labor accessible chromatin regions as early as D17, marking regions that gain tPR binding without PRB enrichment at prelabor, indicating PRA binds with JUND at these regions. In addition, putative enhancers that produce eRNA in the prelaboring and laboring mouse myometrium implicate additional transcription factors from the SOX and ETS families, in addition to AP-1, in driving labor associated gene expression. By contrast, PRB with JUND is associated with gene expression at gestational D17 and in postpartum myometrium. PRB without JUND was also observed at H3K27me3 repressed regions apparent at the prelabor and postpartum stages, which are associated with the repression of labor expressed genes. Furthermore, H3K27me3 erosion, particularly at the HOXD locus appears to contribute to gene expression postpartum. Collectively, these findings highlight the importance of finely tuned transcriptional regulation of the myometrial phenotypic transitions during pregnancy, labor and postpartum.

Contrary to earlier findings that report no changes in euchromatin associated histone modifications (H3K27ac, H3K4me3) between the pregnant and laboring myometrium (Shchuka *et al*., 2020; Dotts *et al*., 2023), ATAC-seq revealed distinct patterns of chromatin accessibility associated with the different stages of pregnancy, labor, and postpartum. These differences may be due to the increased sensitivity of ATAC-seq to changes in chromatin compaction with bp resolution (Bentsen *et al*., 2020). Indeed, regions that do not exhibit changes in bulk chromatin accessibility nevertheless show differences in motif accessibility. We observed that changes in chromatin accessibility did not correspond to alterations in repressive histone marks, which is consistent with prior integration of ATAC-seq and histone ChIP-seq data (Scott *et al*., 2016; Bysani *et al*., 2019). However, the observation that regions displaying H3K27me3 dynamics did not overlap most of the ATAC-seq regions at any studied time point, indicate that H3K27me3 is not involved in repressing enhancers for later activation in the myometrium. For instance, during embryonic stem cell differentiation, poised enhancers play roles in coordinating the initial processes of lineage specification and are characterized by the co-occurrence of H3K4me1 and H3K27me3 (Rada-Iglesias *et al*., 2011).

Analysis of chromatin accessibility changes from labor to postpartum highlighted a large reduction of open chromatin regions following term delivery. Such drastic changes in the active chromatin landscape may be necessary for success of the postpartum uterine involution process, which is important for returning the myometrium to a non-pregnant state. In addition to a massive shutting down of chromatin accessibility for labor expressed genes, D4PP was marked by the opening of new regions with HOX motif enrichment and are largely associated with postpartum expressed genes. HOX factors are known to have pioneering capabilities, allowing them to engage closed chromatin and promote chromatin accessibility (Daftary and Taylor, 2006), a mechanism which has been supported through several studies in the context of limb development in *Drosophila* (reviewed in Cain and Gebelein, 2021). In the context of the myometrium, HOXA10 and HOXA11 have been implicated as a suppressor of labor-associated genes including *IL-8*, *IL-6*, *PTGS2*, and *GJA1* (Liu *et al*., 2015; Li *et al*., 2018). Interestingly, cellular remodeling in uterine fibroids was associated with increased *HOXA13* expression and with changes in ECM-related genes (collagens and MMPs), also upregulated postpartum (George *et al*., 2019). HOXD genes, by contrast, have not been studied in the context of the myometrium and our findings are the first to report upregulation of these genes linked with H3K27me3 erosion postpartum. By day 4 postpartum, though not fully involuted, the uterus regains functional capacity to support embryo implantation, demonstrating the remarkable efficiency of postpartum uterine recovery in mice. The dynamic chromatin footprints and changes in gene expression we identified support a crucial and multifaceted role for HOX family transcription factors in the regulation of genes expressed during postpartum uterine involution.

Intriguingly, accessible chromatin regions at the prelabor and labor stages are associated with JUND and total PR binding but showed no enrichment of PRB, suggesting mainly PRA is bound at these regions. This could be driven by the switch in myometrial PR isoform expression, with PRB predominance observed during gestation, and a PRA increase before labor initiation, elevating the PRA:PRB ratio (Reviewed in Khader *et al*., 2021). In addition, we observed an abundance of AP-1 motifs in the total PR bound regions at prelabor and labor suggesting that PRA association at this time is highly co-occurrent with AP-1 motifs. It has been reported that PRB maintains myometrial quiescence by repressing contraction associated gene expression throughout gestation (Pieber *et al*., 2001; Mesiano *et al*., 2002; Merlino *et al*., 2007; Chai *et al*., 2012; Tan *et al*., 2012; Nadeem *et al*., 2017), however, we observed JUND/PRB co-binding at gestational D17 accessible chromatin regions associated with gene expression on D17, suggesting a more prominent role in gene activation for PRB. We also identified an association of PRB with regions of H3K27me3 repression and these were associated with repressed genes, although predominantly those expressed at postpartum, revealing a new role for PRB in repressing the postpartum stage associated transcriptome during pregnancy. The differential binding of PRB at specific chromatin regions underscores the different roles of this factor in transcriptional regulation of the pregnant and postpartum but not the laboring myometrium where PRB was largely absent.

The analysis of non-coding eRNAs produced specifically during the prelabor and labor stages revealed enhancer candidates, with motif enrichment analysis indicating the involvement of the AP-1, SOX, and ETS transcription factor families. The high occurrence of AP-1 motifs in active chromatin regions of term myometrium supports a genome wide role for AP-1 factors in regulating expression of genes associated with active labor contractions (Mitchell and Lye, 2002, 2005; Khanjani *et al*., 2012; Nadeem *et al*., 2018; Peng *et al*., 2018; Shchuka *et al*., 2020). The enrichment of ETS motifs in contractility-associated enhancers is consistent with earlier findings of enriched ETS motifs in H3K27ac regions obtained from the laboring human myometrium (Dotts *et al*., 2023). One ETS family member, *Elf3*, exhibits increased expression at prelabor and labor compared to both D17 and D4PP, aligning with prior findings using laboring myometrium tissues (Bethin *et al*., 2003; Shchuka *et al*., 2020). *Elf3* has also been shown to have an effect on labor-associated gene promoters in collaboration with AP-1 factors in different cell types, including myometrial cells (Otero *et al*., 2012, 2017; Shchuka *et al*., 2023). Similarly, SOX factor motifs were previously found to be enriched in H3K4me3 marked regions of the laboring human myometrium (Dotts *et al*., 2023) and had a demonstrated regulatory role, alongside AP-1 dimers, on labor-driving gene promoters including *Fos, Mmp11, Ptgs2,* and *Cald1* (Jang *et al*., 2015; Ghouili *et al*., 2018; Najih *et al*., 2022; Khader *et al*., 2024). Their role is further supported by RNA-seq analysis from this study which displays differential expression of SOX family genes throughout gestation. In particular, *Sox4*, *Sox6*, *Sox7*, and *Sox11*, are upregulated at prelabor compared to D17; *Sox6* and *Sox11* are upregulated at prelabor compared to labor; and *Sox7* and *Sox9* are increased at labor compared to D4PP. SOX4 protein levels were elevated in laboring mouse and human myometrium relative to non-laboring tissues (Bethin *et al*., 2003; Fan *et al*., 2020; Khader *et al*., 2024). SOX factors have also been shown to interact with ETS family members in different cellular contexts, such as the repressive activity of ELF3 through direct interactions with SOX9 to inhibit the expression of *COL2A1* (Otero *et al*., 2017). These findings suggest a complex relationship between AP-1, SOX, and ETS transcription factor family members in regulating uterine contractility by modulation of contraction associated gene expression to control the switch in myometrial phenotypes and labor onset.

It is worth noting that the data presented in this study are gathered from bulk myometrial tissue and therefore are not able to reveal any of the additional complexities that arise from changes in the complement of cells present in the myometrium. Of particular interest would be the infiltration of immune cells that occurs at term and plays a role in labor onset (Shynlova *et al*., 2013b, 2013a, 2021; Srikhajon *et al*., 2014), although these cells represent a small number of the total cell count (<2%) (Sweeney *et al*., 2013) and are therefore not expected to skew the ATAC-seq or ChIP-seq data presented. Although myometrial smooth cells have been estimated to make up more than 65% of the myometrial layer of the uterus (Sweeney *et al*., 2013), fibroblasts are also present within this layer and have been recently implicated in controlling the timing of labor onset through chromatin regulatory mechanisms (McIntyre *et al*., 2025).

## Conclusions

This study is the largest collection of genome-wide data profiling the temporal transcriptomic, epigenomic, and transcription factor binding changes that occur in the mouse myometrium throughout gestation, labor and postpartum. Our data (1) cemented the role of AP-1 factors as genome-wide regulators of myometrial gene expression, (2) implicated ETS and SOX factors in the contractility-associate transcriptional regulation of term labor, and (3) identified HOX factors as regulators of postpartum uterine involution. Gaining a clearer insight into the key transcription factors regulating different myometrial phenotypes enhances our understanding of the complexity of the molecular mechanisms governing reproduction. This knowledge can potentially aid in identifying novel drug targets for prevention of pregnancy complications such as preterm labor or postpartum hemorrhage.

## Methods

### Ethics statement

This study was carried out in accordance with the protocol approved by the Research Ethics Board, Sinai Health System REB# 02-0061A and REB# 18-0168A. All subjects donating myometrial biopsies for research gave written informed consent in accordance with the Declaration of Helsinki. All research using animal tissues was performed in a class II certified laboratory by qualified staff trained in biological and chemical safety protocols and in accordance with Health Canada guidelines and regulations.

### Animal model tissue collection

This study was carried out in accordance with standards set out by the Canadian Council on Animal Care (CCAC). All animal experiments were reviewed and approved by the Animal Care Committee of The Centre for Phenogenomics and the Animal Use Protocol (AUP# 21-0164H). Virgin outbred CD-1 mice used in these experiments were purchased from Harlan Laboratories (http://www.harlan.com/). All animals in this AUP were maintained and used in accordance with the current recommendations of CCAC, the requirements under the Animals for Research Act, RSO 1990, and The Centre for Phenogenomics Committee Policies and Guidelines. All mice were housed under specific pathogen–free conditions on a 12L:12D cycle and were administered food and water ad libitum. Female CD-1 mice were mated overnight with males, and day 0.5 of gestation was designated upon the observation of a vaginal plug. Pregnant mice were maintained until the appropriate gestational time point. The average time of delivery under these conditions was on day 19.5 and labor was determined based on the delivery of at least 1 pup. Animals were sacrificed and the myometrium samples were obtained on gestational days 15, 17, prelabor (PL; day 19), during term labor (day 19.5), one day post-partum (day 20) and four days postpartum (day 23). The myometrium tissue was removed using a cutting/scraping method to remove the decidua basalis and decidua parietalis, frozen in liquid nitrogen, and stored at -80°C. All mice were stored at the Toronto Centre for Phenogenomics (TCP). All animal experiments were conducted by the Lye Lab with approval by the TCP Animal Care Committee.

### Assay for transposase accessible chromatin paired with sequencing (ATAC-seq) data analysis

Raw fastq files were analyzed using the ENCODE ATAC-seq pipeline as established by the Kundaje Lab (“ENCODE-DCC/atac-seq-pipeline,” 2024). Briefly, raw files were assessed for quality using FASTQC and aligned to the mm10 genome using Bowtie2. MACS2 (Zhang *et al*., 2008) was used for peak-calling and significantly conserved peaks in biological replicates were filtered using Irreproducible Discovery Rate (IDR) (Li *et al*., 2011). These conserved IDR peaks were retained for downstream analyses. Bam files of all replicates per sample group were merged using Samtools *merge* and used to create bedgraphs that were normalized to reads per million mapped reads (RPM), via bamCoverage from Deeptools. (Ramírez *et al*., 2016) computeMatrix from DeepTools was used to generate TSS enrichment plots of peak enrichment at transcription start site (TSS). Peak annotation was conducted using Homer, *annotatePeaks.pl* (Heinz *et al*., 2010). Footprinting analysis was done via TOBIAS (Bentsen *et al*., 2020) and results were visualized through volcano plots using EnhancedVolcano (Blighe *et al*., 2023). The diffBind package (Ross-Innes *et al*., 2012; Stark and Brown, 2023) was used to conduct the differentially accessible enrichment analysis between different time points (log_2_FC of ≤ 1 and ≥ 1; adjusted *p*-value of < 0.05], through which the Venn diagrams, Volcano Plot, and heatmaps (dba.plotVenn, dba.plotVolcano, dba.plotHeatmap, respectively) were generated. ChIPSeeker was used to annotate genes and genomic annotation and create plots (Yu *et al*., 2015).

Sequencing data files were submitted to the GEO repository (GSE158675).

### RNA extraction

Total RNA was extracted from frozen, crushed myometrium tissue samples, obtained from mice at D17, PL (d19), LAB (D19.5), D4PP (D23) (n = 3 each), using a Trizol/Chloroform extraction protocol. RNA samples were column-purified using RNeasy Mini Kit (Qiagen), treated with DNase I (Qiagen) to remove genomic DNA contamination, and subsequently subjected to standard Illumina Stranded Total RNA Prep Ligation with Ribo-Zero Plus protocol for paired-end sequencing.

### RNA-seq data analysis

Raw data of FASTQ format were quality assessed and trimmed using fastp (Chen *et al*., 2018) followed by mapping to the mm10 genome using STAR (Dobin *et al*., 2013). Reads were quantified using featureCounts (exon reads) (Liao *et al*., 2014) and intron reads were quantified using SeqMonk’s active transcription quantitation pipeline. Alternative transcript counts were summed together for every gene. Intron reads were then imported into DESeq2 [for differential expression analysis (Love *et al*., 2014). Genes with a |log_2_FC| ≥ 1.5 and adjusted *p*-value ≤ 0.01 were considered significantly different. DESeq2 was used to assess the differential expression and data were plotted using the EnhancedVolcano (Blighe *et al*., 2023) and the ComplexHeatmap (Gu *et al*., 2016; Gu, 2022) package in R, for heatmaps and volcano plots, respectively. Enrichment results were sorted by the adjusted p-value and grouped by the biological process. DeepTools (bamCoverage) (Ramírez *et al*., 2016) was used to generate bedgraphs which were visualized on UCSC. Sequencing data files were submitted to the GEO repository (GSE217817).

### Chromatin immunoprecipitation paired with sequencing (ChIP-seq)

Chromatin immunoprecipitation was performed as described previously (Taylor et al., 2022) with minor modifications. Flash frozen mouse myometrium tissues (100 mg) obtained from Day 17, PL, LAB mice were crushed and cross-linked using 1% formaldehyde for ten minutes, followed by quenching through adding glycine to a final concentration of 125 mM for 15 min with rotation. For JUND ChIPs, flash-frozen tissues were subjected to a secondary cross-linker, disuccinimidyl glutarate (DSG), for 30 mins at room temperature, prior to formaldehyde fixation using 1% formaldehyde for ten minutes. Cell pellets were washed twice with ice-cold PBS followed by resuspension in ice-cold lysis buffer 1 (50 mM HEPES-KOH, 140 mM NaCl, 1 mM EDTA, 10% glycerol, 0.5% NP-40, 0.25% Triton X-100) for 10 min at 4°C with rotation.

Lysates were centrifuged at 2000g for 5 min at 4°C before resuspending in ice-cold lysis buffer 2 (10 mM TrisHCl, 200 mM NaCl, 1 mM EDTA, 0.5 mM EGTA) for 10 min at 4°C. Nuclei were pelleted at 2000g for 5 min at 4°C before resuspending in ice-cold lysis buffer 3 (10 mM Tris-HCl, 100 mM NaCl, 1 mM EDTA, 0.5 mM EGTA, 0.1% Na-deoxycholate, 0.5% N-laurylsarcosine). Sonication was performed using a probe sonicator at 20 Amps (15 sec on/30 sec off) for 2 min at 4°C. After cell lysis and sonication, Triton X-100 was added to the sonicated lysate to precipitate any debris. Fifty microliters of cell lysate were retained as whole-cell extract (WCE), and the remaining lysate was split between two different immunoprecipitations (in the case of PRB and H3K27me3 ChIPs). Chromatin lysates were incubated overnight with 5 μg of relevant antibodies at 4°C with rotation. Antibodies used for ChIP experiments were antibodies targeting the progesterone receptor, total PR (ThermoFisher; MA5-14505), PRB (abcam; ab2765), JUND (abcam; ab181615), and H3K27me3 (abcam; ab192985).

Protein A magnetic Dynabeads (80 uL, Invitrogen) were washed thrice with 1% BSA/PBS before adding to the antibody-bound chromatin lysates and incubated overnight at 4°C with rotation. Lysates were washed four times with 1mL of room temperature RIPA buffer and then with a TBS buffer. Antibody-bound lysates and WCE were eluted in an SDS-based elution buffer (50 mM Tris-HCl, 10 mM EDTA, 1% SDS) for 30 min at 65°C before addition of proteinase K and overnight reverse cross-linking at 65°C with shaking at 900 RPM. ChIP DNA was purified using the phenol/ chloroform method and eluted in 65 uL of nuclease-free water. Aliquots from the sonicated samples were run on a 1% agarose gel to confirm that chromatin was sheared to a range of 300–1000 bp in size. ChIP samples were submitted for paired-end 150bp read sequencing using standard Illumina HiSeq 2500 protocols.

### ChIP-seq data analysis

Raw fastq reads were quality assessed and trimmed using fastp and mapped to the mm10 reference genome using STAR (Dobin *et al*., 2013). ChIP-seq peaks were called, relative to the input, for each individual replicate using MACS2 (Zhang *et al*., 2008) with default parameters (*p* < 0.01) (broadpeaks for H3K27me3 and narrowpeaks for transcription factor ChIPs). Significantly conserved peaks between biological replicates were combined using irreproducible discovery rate (IDR) (Li *et al*., 2011). Differential peak analysis was performed using the DiffBind package (Ross-Innes *et al*., 2012; Stark and Brown, 2023). Peaks with an |log_2_FC| ≥ 1 and adjusted *p*-value < 0.05 were considered significantly different between various time point comparisons. Fischer’s test and Jaccard index was conducted via bedtools (Quinlan and Hall, 2010). ChIPSeeker was used to annotate genes and genomic annotation and create plots (Yu *et al*., 2015). Sequencing data files were submitted to the GEO repository (GSE240819 and GSE291680). Enrichment plots were created using computeMatrix from deepTools.

## Acknowledgements

We would like to thank the staff at the Princess Margaret Genomics Centre for conducting the ATAC-seq with the mouse myometrium tissues. Thank you to the members of the Mitchell Lab, especially Dr. Ian C. Tobias for insightful discussions regarding bioinformatic analyses. Figure schematics in figures 1A and supplemental figure 3A, were created with Biorender.com. This research was supported by funding from the Canadian Institutes of Health Research (https://cihr-irsc.gc.ca/), PJT-173252 held by JAM and OS. The funders had no role in the study design, data collection and analysis, decision to publish, or preparation of the manuscript.

## References

1. Arrowsmith S, Wray S. Oxytocin: Its Mechanism of Action and Receptor Signalling in the Myometrium. J Neuroendocrinology 2014;26:356–369.

2. Bentsen M, Goymann P, Schultheis H, Klee K, Petrova A, Wiegandt R, Fust A, Preussner J, Kuenne C, Braun T, et al. ATAC-seq footprinting unravels kinetics of transcription factor binding during zygotic genome activation. Nat Commun 2020;11:4267. Nature Publishing Group.

3. Bethin KE, Nagai Y, Sladek R, Asada M, Sadovsky Y, Hudson TJ, Muglia LJ. Microarray Analysis of Uterine Gene Expression in Mouse and Human Pregnancy. Molecular Endocrinology 2003;17:1454–1469.

4. Blighe K, Rana S, Turkes E, Ostendorf B, Grioni A, Lewis M. EnhancedVolcano: Publication-ready volcano plots with enhanced colouring and labeling. Bioconductor [Internet] 2023;Available from: http://bioconductor.org/packages/EnhancedVolcano/.

5. Bollopragada S, Youssef R, Jordan F, Greer I, Norman J, Nelson S. Term labor is associated with a core inflammatory response in human fetal membranes, myometrium, and cervix. American Journal of Obstetrics & Gynecology 2009;200:104.e1–104.e11. Elsevier.

6. Bose DA, Donahue G, Reinberg D, Shiekhattar R, Bonasio R, Berger SL. RNA Binding to CBP Stimulates Histone Acetylation and Transcription. Cell 2017;168:135–149.e22.

7. Bysani M, Agren R, Davegårdh C, Volkov P, Rönn T, Unneberg P, Bacos K, Ling C. ATAC-seq reveals alterations in open chromatin in pancreatic islets from subjects with type 2 diabetes. Sci Rep 2019;9:7785.

8. Cain B, Gebelein B. Mechanisms Underlying Hox-Mediated Transcriptional Outcomes. Front Cell Dev Biol 2021;9:787339.

9. Cao R, Wang L, Wang H, Xia L, Erdjument-Bromage H, Tempst P, Jones RS, Zhang Y. Role of Histone H3 Lysine 27 Methylation in Polycomb-Group Silencing. Science 2002;298:1039–1043. American Association for the Advancement of Science.

10. Chai SY, Smith R, Zakar T, Mitchell C, Madsen G. Term myometrium is characterized by increased activating epigenetic modifications at the progesterone receptor-A promoter. Mol Hum Reprod 2012;18:401–409.

11. Chan Y-W, Berg HA van den, Moore JD, Quenby S, Blanks AM. Assessment of myometrial transcriptome changes associated with spontaneous human labour by high-throughput RNA-seq. Exp Physiol 2014;99:510–524.

12. Chen S, Zhou Y, Chen Y, Gu J. fastp: an ultra-fast all-in-one FASTQ preprocessor. Bioinformatics 2018;34:i884–i890.

13. Creyghton MP, Cheng AW, Welstead GG, Kooistra T, Carey BW, Steine EJ, Hanna J, Lodato MA, Frampton GM, Sharp PA, et al. Histone H3K27ac separates active from poised enhancers and predicts developmental state. Proceedings of the National Academy of Sciences 2010;107:21931–21936. Proceedings of the National Academy of Sciences.

14. Daftary GS, Taylor HS. Endocrine Regulation of HOX Genes. Endocrine Reviews 2006;27:331–355.

15. Dobin A, Davis CA, Schlesinger F, Drenkow J, Zaleski C, Jha S, Batut P, Chaisson M, Gingeras TR. STAR: ultrafast universal RNA-seq aligner. Bioinformatics 2013;29:15–21.

16. Dong X, Yu C, Shynlova O, Challis JRG, Rennie PS, Lye SJ. p54nrb Is a Transcriptional Corepressor of the Progesterone Receptor that Modulates Transcription of the Labor-Associated Gene, Connexin 43 (Gja1). Mol Endocrinol 2009;23:1147–1160. Oxford Academic.

17. Dotts AJ, Reiman D, Yin P, Kujawa S, Grobman WA, Dai Y, Bulun SE. In Vivo Genome-Wide PGR Binding in Pregnant Human Myometrium Identifies Potential Regulators of Labor. Reprod Sci 2023;30:544–559. Thousand Oaks, Calif.

18. ENCODE-DCC/atac-seq-pipeline 2024; ENCODE DCCAvailable from: https://github.com/ENCODE-DCC/atac-seq-pipeline.

19. Fan Y, Hou W, Xing Y, Zhang L, Zhou C, Gui J, Xu P, Wang A, Fan X, Zeng X, et al. Peptidomics analysis of myometrium tissues in term labor compared with term nonlabor. Journal of Cellular Biochemistry 2020;121:1890–1900.

20. Fischle W, Wang Y, Jacobs SA, Kim Y, Allis CD, Khorasanizadeh S. Molecular basis for the discrimination of repressive methyl-lysine marks in histone H3 by Polycomb and HP1 chromodomains. Genes Dev 2003;17:1870–1881.

21. George JW, Fan H, Johnson B, Carpenter TJ, Foy KK, Chatterjee A, Patterson AL, Koeman J, Adams M, Madaj ZB, et al. Integrated Epigenome, Exome, and Transcriptome Analyses Reveal Molecular Subtypes and Homeotic Transformation in Uterine Fibroids. Cell Rep 2019;29:4069–4085.e6.

22. Ghouili F, Roumaud P, Martin LJ. Gja1 expression is regulated by cooperation between SOX8/SOX9 and cJUN transcription factors in TM4 and 15P-1 Sertoli cell lines. Molecular Reproduction and Development 2018;85:875–886.

23. Gu Z. Complex heatmap visualization. iMeta 2022;1:e43.

24. Gu Z, Eils R, Schlesner M. Complex heatmaps reveal patterns and correlations in multidimensional genomic data. Bioinformatics 2016;32:2847–2849.

25. Hackwell ECR, Ladyman SR, Brown RSE, Grattan DR. Mechanisms of Lactation-induced Infertility in Female Mice. Endocrinology 2023;164:bqad049.

26. Heintzman ND, Stuart RK, Hon G, Fu Y, Ching CW, Hawkins RD, Barrera LO, Van Calcar S, Qu C, Ching KA, et al. Distinct and predictive chromatin signatures of transcriptional promoters and enhancers in the human genome. Nat Genet 2007;39:311–318. Nature Publishing Group.

27. Heinz S, Benner C, Spann N, Bertolino E, Lin YC, Laslo P, Cheng JX, Murre C, Singh H, Glass CK. Simple Combinations of Lineage-Determining Transcription Factors Prime cis-Regulatory Elements Required for Macrophage and B Cell Identities. Molecular Cell 2010;38:576–589. Elsevier.

28. Hsu K-F, Pan H-A, Hsu Y-Y, Wu C-M, Chung W-J, Huang S-C. Enhanced myometrial autophagy in postpartum uterine involution. Taiwan J Obstet Gynecol 2014;53:293–302.

29. Ishihara K, Oshimura M, Nakao M. CTCF-Dependent Chromatin Insulator Is Linked to Epigenetic Remodeling. Molecular Cell 2006;23:733–742. Elsevier.

30. Jang S-M, Kim J-W, Kim C-H, An J-H, Johnson A, Song PI, Rhee S, Choi K-H. KAT5-mediated SOX4 acetylation orchestrates chromatin remodeling during myoblast differentiation. Cell Death Dis 2015;6:e1857–e1857. Nature Publishing Group.

31. Kaikkonen MU, Spann NJ, Heinz S, Romanoski CE, Allison KA, Stender JD, Chun HB, Tough DF, Prinjha RK, Benner C, et al. Remodeling of the Enhancer Landscape during Macrophage Activation Is Coupled to Enhancer Transcription. Molecular Cell 2013;51:310–325.

32. Khader N, Dorogin A, Shynlova O, Mitchell JA. JUND plays a genome-wide role in the quiescent to contractile switch in the pregnant human myometrium. PLoS Genet 2025;21:e1011261.

33. Khader N, Shchuka VM, Dorogin A, Shynlova O, Mitchell JA. SOX4 exerts contrasting regulatory effects on labor-associated gene promoters in myometrial cells. PLoS One 2024;19:e0297847.

34. Khader N, Shchuka VM, Shynlova O, Mitchell JA. Transcriptional control of parturition: insights from gene regulation studies in the myometrium. Mol Hum Reprod 2021;27:gaab024.

35. Khanjani S, Terzidou V, Johnson MR, Bennett PR. NFκB and AP-1 Drive Human Myometrial IL8 Expression. Mediators of Inflammation [Internet] 2012;2012:e504952. Hindawi.

36. Kimura T, Takemura M, Nomura S, Nobunaga T, Kubota Y, Inoue T, Hashimoto K, Kumazawa I, Ito Y, Ohashi K, et al. Expression of oxytocin receptor in human pregnant myometrium. Endocrinology 1996;137:780–785.

37. Li Q, Brown JB, Huang H, Bickel PJ. Measuring reproducibility of high-throughput experiments. The Annals of Applied Statistics 2011;5:1752–1779. Institute of Mathematical Statistics.

38. Li Y, Li H, Xie N, Chen R, Lee AR, Slater D, Lye S, Dong X. HoxA10 and HoxA11 Regulate the Expression of Contraction-Associated Proteins and Contribute to Regionalized Myometrium Phenotypes in Women. Reprod Sci 2018;25:44–50.

39. Liang M, Soomro A, Tasneem S, Abatti LE, Alizada A, Yuan X, Uusküla-Reimand L, Antounians L, Alvi SA, Paterson AD, et al. Enhancer-gene rewiring in the pathogenesis of Quebec platelet disorder. Blood 2020;136:2679–2690.

40. Liao Y, Smyth GK, Shi W. featureCounts: an efficient general purpose program for assigning sequence reads to genomic features. Bioinformatics 2014;30:923–930.

41. Liu L, Li H, Dargahi D, Shynlova O, Slater D, Jones SJM, Lye SJ, Dong X. HoxA13 Regulates Phenotype Regionalization of Human Pregnant Myometrium. J Clin Endocrinol Metab 2015;100:E1512–1522.

42. Love MI, Huber W, Anders S. Moderated estimation of fold change and dispersion for RNA-seq data with DESeq2. Genome Biology 2014;15:550.

43. Lye SJ, Mitchell J, Nashman N, Oldenhof A, Ou R, Shynlova O, Langille L. Role of mechanical signals in the onset of term and preterm labor. Front Horm Res 2001;27:165–178.

44. Madsen JGS, Schmidt SF, Larsen BD, Loft A, Nielsen R, Mandrup S. iRNA-seq: computational method for genome-wide assessment of acute transcriptional regulation from total RNA-seq data. Nucleic Acids Res 2015;43:e40.

45. McIntyre TI, Valdez O, Kochhar NP, Davidson B, Samad B, Qiu L, Hu K, Combes AJ, Erlebacher A. KDM6B-dependent epigenetic programming of uterine fibroblasts in early pregnancy regulates parturition timing in mice. Cell 2025;188:1265–1279.e18.

46. Merlino AA, Welsh TN, Tan H, Yi LJ, Cannon V, Mercer BM, Mesiano S. Nuclear progesterone receptors in the human pregnancy myometrium: evidence that parturition involves functional progesterone withdrawal mediated by increased expression of progesterone receptor-A. J Clin Endocrinol Metab 2007;92:1927–1933.

47. Mesiano S, Chan E-C, Fitter JT, Kwek K, Yeo G, Smith R. Progesterone withdrawal and estrogen activation in human parturition are coordinated by progesterone receptor A expression in the myometrium. J Clin Endocrinol Metab 2002;87:2924–2930.

48. Mitchell JA, Lye SJ. Differential Expression of Activator Protein-1 Transcription Factors in Pregnant Rat Myometrium. Biol Reprod 2002;67:240–246. Oxford Academic.

49. Mitchell JA, Lye SJ. Differential Activation of the Connexin 43 Promoter by Dimers of Activator Protein-1 Transcription Factors in Myometrial Cells. 2005;146:2048–2054.

50. Mitchell JA, Shynlova O, Langille BL, Lye SJ. Mechanical stretch and progesterone differentially regulate activator protein-1 transcription factors in primary rat myometrial smooth muscle cells. American Journal of Physiology-Endocrinology and Metabolism 2004;287:E439–E445.

51. Mittal P, Romero R, Tarca AL, Gonzalez J, Draghici S, Xu Y, Dong Z, Nhan-Chang C-L, Chaiworapongsa T, Lye S, et al. Characterization of the myometrial transcriptome and biological pathways of spontaneous human labor at term. J Perinat Med 2010;38:617–643.

52. Moorthy SD, Davidson S, Shchuka VM, Singh G, Malek-Gilani N, Langroudi L, Martchenko A, So V, Macpherson NN, Mitchell JA. Enhancers and super-enhancers have an equivalent regulatory role in embryonic stem cells through regulation of single or multiple genes. Genome Res 2017;27:246–258.

53. Mousavi K, Zare H, Dell’Orso S, Grontved L, Gutierrez-Cruz G, Derfoul A, Hager GL, Sartorelli V. eRNAs Promote Transcription by Establishing Chromatin Accessibility at Defined Genomic Loci. Molecular Cell 2013;51:606–617.

54. Nadeem L, Farine T, Dorogin A, Matysiak-Zablocki E, Shynlova O, Lye S. Differential expression of myometrial AP-1 proteins during gestation and labour. J Cell Mol Med 2018;22:452–471.

55. Nadeem L, Shynlova O, Matysiak-Zablocki E, Mesiano S, Dong X, Lye S. Molecular evidence of functional progesterone withdrawal in human myometrium. Nature Communications 2016;7:11565. Nature Publishing Group.

56. Nadeem L, Shynlova O, Mesiano S, Lye S. Progesterone Via its Type-A Receptor Promotes Myometrial Gap Junction Coupling. Sci Rep 2017;7:13357.

57. Najih M, Basque A, Nguyen HT, Diawara M, Martin LJ. Members of the AP-1 Family of Transcription Factors Regulate the Expression of Gja1 in Mouse GC-1 Spermatogonial Cells. Applied Sciences 2022;12:1408. Multidisciplinary Digital Publishing Institute.

58. Nishinaka K, Fukuda Y. Changes in extracellular matrix materials in the uterine myometrium of rats during pregnancy and postparturition. Acta Pathol Jpn 1991;41:122–132.

59. Otero M, Peng H, El Hachem K, Culley KL, Wondimu EB, Quinn J, Asahara H, Tsuchimochi K, Hashimoto K, Goldring MB. ELF3 modulates type II collagen gene (COL2A1) transcription in chondrocytes by Inhibiting SOX9-CBP/p300-driven histone acetyltransferase activity. Connect Tissue Res 2017;58:15–26.

60. Otero M, Plumb DA, Tsuchimochi K, Dragomir CL, Hashimoto K, Peng H, Olivotto E, Bevilacqua M, Tan L, Yang Z, et al. E74-like Factor 3 (ELF3) Impacts on Matrix Metalloproteinase 13 (MMP13) Transcriptional Control in Articular Chondrocytes under Proinflammatory Stress. J Biol Chem 2012;287:3559–3572.

61. Ou C-W, Chen Z-Q, Qi S, Lye SJ. Increased Expression of the Rat Myometrial Oxytocin Receptor Messenger Ribonucleic Acid during Labor Requires Both Mechanical and Hormonal Signals1. Biology of Reproduction 1998;59:1055–1061.

62. Ou CW, Orsino A, Lye SJ. Expression of connexin-43 and connexin-26 in the rat myometrium during pregnancy and labor is differentially regulated by mechanical and hormonal signals. Endocrinology 1997;138:5398–5407.

63. Peng Q, Liu Y, Dong M, Xu F, Huang J, Chen J, Li X, Zhang J, Zhang W. Interaction between NF-κB and AP-1 and their intracellular localization at labor in human late pregnant myometrial cells in vivo and in vitro. Medicine (Baltimore) [Internet] 2018;97:.

64. Phillips JE, Corces VG. CTCF: Master Weaver of the Genome. Cell 2009;137:1194–1211. Elsevier.

65. Pieber D, Allport VC, Hills F, Johnson M, Bennett PR. Interactions between progesterone receptor isoforms in myometrial cells in human labour. Mol Hum Reprod 2001;7:875–879.

66. Quinlan AR, Hall IM. BEDTools: a flexible suite of utilities for comparing genomic features. Bioinformatics 2010;26:841–842.

67. Rada-Iglesias A, Bajpai R, Swigut T, Brugmann SA, Flynn RA, Wysocka J. A unique chromatin signature uncovers early developmental enhancers in humans. Nature 2011;470:279–283.

68. Ramírez F, Ryan DP, Grüning B, Bhardwaj V, Kilpert F, Richter AS, Heyne S, Dündar F, Manke T. deepTools2: a next generation web server for deep-sequencing data analysis. Nucleic Acids Research 2016;44:W160–W165.

69. Ross-Innes CS, Stark R, Teschendorff AE, Holmes KA, Ali HR, Dunning MJ, Brown GD, Gojis O, Ellis IO, Green AR, et al. Differential oestrogen receptor binding is associated with clinical outcome in breast cancer. Nature 2012;481:389–393. Nature Publishing Group.

70. Rubel CA, Lanz RB, Kommagani R, Franco HL, Lydon JP, DeMayo FJ. Research Resource: Genome-Wide Profiling of Progesterone Receptor Binding in the Mouse Uterus. Mol Endocrinol 2012;26:1428–1442.

71. Sartorelli V, Lauberth SM. Enhancer RNAs are an important regulatory layer of the epigenome. Nat Struct Mol Biol 2020;27:521–528. Nature Publishing Group.

72. Scott MT, Korfi K, Saffrey P, Hopcroft LEM, Kinstrie R, Pellicano F, Guenther C, Gallipoli P, Cruz M, Dunn K, et al. Epigenetic Reprogramming Sensitizes CML Stem Cells to Combined EZH2 and Tyrosine Kinase Inhibition. Cancer Discov 2016;6:1248–1257.

73. Shchuka VM, Abatti LE, Hou H, Khader N, Dorogin A, Wilson MD, Shynlova O, Mitchell JA. The pregnant myometrium is epigenetically activated at contractility-driving gene loci prior to the onset of labor in mice. PLoS Biol 2020;18:e3000710.

74. Shchuka VM, Khader N, Dorogin A, Shynlova O, Mitchell JA. MYB and ELF3 differentially modulate labor-inducing gene expression in myometrial cells. PLOS ONE 2023;18:e0271081. Public Library of Science.

75. Shynlova O, Boros-Rausch A, Farine T, Adams Waldorf KM, Dunk C, Lye SJ. Decidual Inflammation Drives Chemokine-Mediated Immune Infiltration Contributing to Term Labor. J Immunol 2021;207:2015–2026. Baltimore, Md. : 1950.

76. Shynlova O, Lee Y-H, Srikhajon K, Lye SJ. Physiologic uterine inflammation and labor onset: integration of endocrine and mechanical signals. Reprod Sci 2013a;20:154–167. Thousand Oaks, Calif.

77. Shynlova O, Mitchell JA, Tsampalieros A, Langille BL, Lye SJ. Progesterone and gravidity differentially regulate expression of extracellular matrix components in the pregnant rat myometrium. Biol Reprod 2004;70:986–992.

78. Shynlova O, Nedd-Roderique T, Li Y, Dorogin A, Lye SJ. Myometrial immune cells contribute to term parturition, preterm labour and post-partum involution in mice. J Cell Mol Med 2013b;17:90–102.

79. Shynlova O, Tsui P, Jaffer S, Lye SJ. Integration of endocrine and mechanical signals in the regulation of myometrial functions during pregnancy and labour. Eur J Obstet Gynecol Reprod Biol 2009;144 **Suppl 1**:S2–10.

80. Singh G, Mullany S, Moorthy SD, Zhang R, Mehdi T, Tian R, Duncan AG, Moses AM, Mitchell JA. A flexible repertoire of transcription factor binding sites and a diversity threshold determines enhancer activity in embryonic stem cells. Genome Res 2021;31:564–575.

81. Srikhajon K, Shynlova O, Preechapornprasert A, Chanrachakul B, Lye S. A new role for monocytes in modulating myometrial inflammation during human labor. Biol Reprod 2014;91:10.

82. Stanfield Z, Lai PF, Lei K, Johnson MR, Blanks AM, Romero R, Chance MR, Mesiano S, Koyutürk M. Myometrial Transcriptional Signatures of Human Parturition. Front Genet 2019;10:185.

83. Stark R, Brown G. DiffBind: Differential binding analysis of ChIP-Seq peak data. 2023;

84. Suarez AC, Gimenez CJ, Russell SR, Wang M, Munson JM, Myers KM, Miller KS, Abramowitch SD, De Vita R. Pregnancy-induced remodeling of the murine reproductive tract: a longitudinal in vivo magnetic resonance imaging study. Sci Rep 2024;14:586.

85. Sweeney EM, Crankshaw DJ, O’Brien Y, Dockery P, Morrison JJ. Stereology of human myometrium in pregnancy: influence of maternal body mass index and age. Am J Obstet Gynecol 2013;208:324.e1-6.

86. Tan H, Yi L, Rote NS, Hurd WW, Mesiano S. Progesterone receptor-A and -B have opposite effects on proinflammatory gene expression in human myometrial cells: implications for progesterone actions in human pregnancy and parturition. J Clin Endocrinol Metab 2012;97:E719–730.

87. Taylor T, Sikorska N, Shchuka VM, Chahar S, Ji C, Macpherson NN, Moorthy SD, Kort MAC de, Mullany S, Khader N, et al. Transcriptional regulation and chromatin architecture maintenance are decoupled functions at the Sox2 locus. Genes Dev 2022;36:699–717.

88. Tobias IC, Abatti LE, Moorthy SD, Mullany S, Taylor T, Khader N, Filice MA, Mitchell JA. Transcriptional enhancers: from prediction to functional assessment on a genome-wide scale. Genome 2021;64:426–448. NRC Research Press.

89. Tsai P-F, Dell’Orso S, Rodriguez J, Vivanco KO, Ko K-D, Jiang K, Juan AH, Sarshad AA, Vian L, Tran M, et al. A Muscle-Specific Enhancer RNA Mediates Cohesin Recruitment and Regulates Transcription In trans. Molecular Cell 2018;71:129–141.e8.

90. Wu S-P, Anderson ML, Wang T, Zhou L, Emery OM, Li X, DeMayo FJ. Dynamic transcriptome, accessible genome, and PGR cistrome profiles in the human myometrium. The FASEB Journal 2020;34:2252–2268.

91. Yu G, Wang L-G, He Q-Y. ChIPseeker: an R/Bioconductor package for ChIP peak annotation, comparison and visualization. Bioinformatics 2015;31:2382–2383.

92. Zenk F, Loeser E, Schiavo R, Kilpert F, Bogdanović O, Iovino N. Germ line-inherited H3K27me3 restricts enhancer function during maternal-to-zygotic transition. Science 2017;357:212–216. New York, N.Y.

93. Zhang Y, Liu T, Meyer CA, Eeckhoute J, Johnson DS, Bernstein BE, Nusbaum C, Myers RM, Brown M, Li W, et al. Model-based Analysis of ChIP-Seq (MACS). Genome Biology 2008;9:R137.

