## Supplemental figures for "Temporal chromatin and transcriptome dynamics driven by JUND and progesterone receptor binding in the pregnant mouse myometrium"

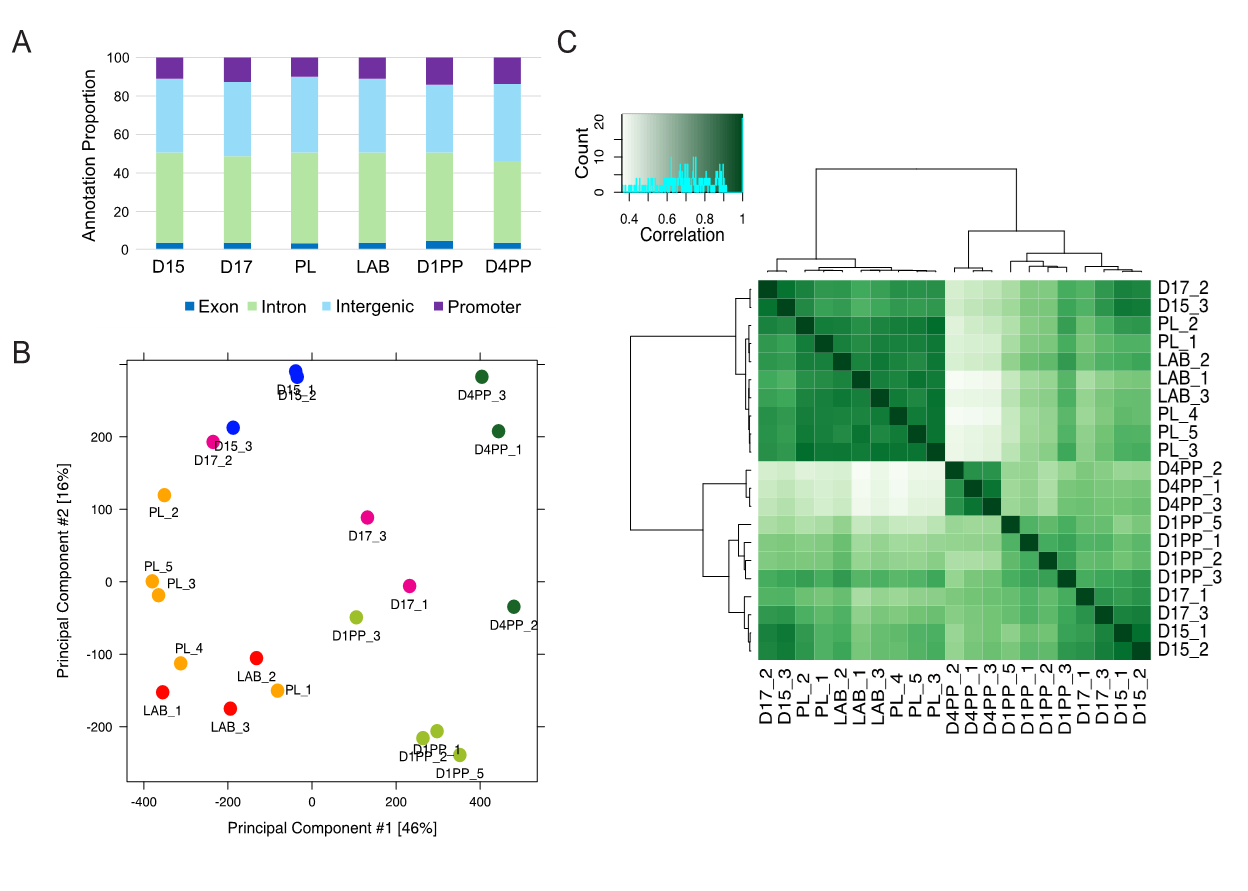

**Supplemental Figure 1: Evaluating chromatin accessibility profiles by ATAC-seq throughout mouse gestation, labor and postpartum.**

A) Functional genomic annotation of peaks found in open chromatin regions at all timepoints. B) PCA clustering shows that ATAC-seq peaks from day 15 and day 17 cluster together and are distinct from those obtained from term (day 19; PL and day 19.5; LAB), and postpartum (day 20; D1PP and day 23; D4PP). C) Hierarchical clustering of ATAC-seq replicates for all timepoints.

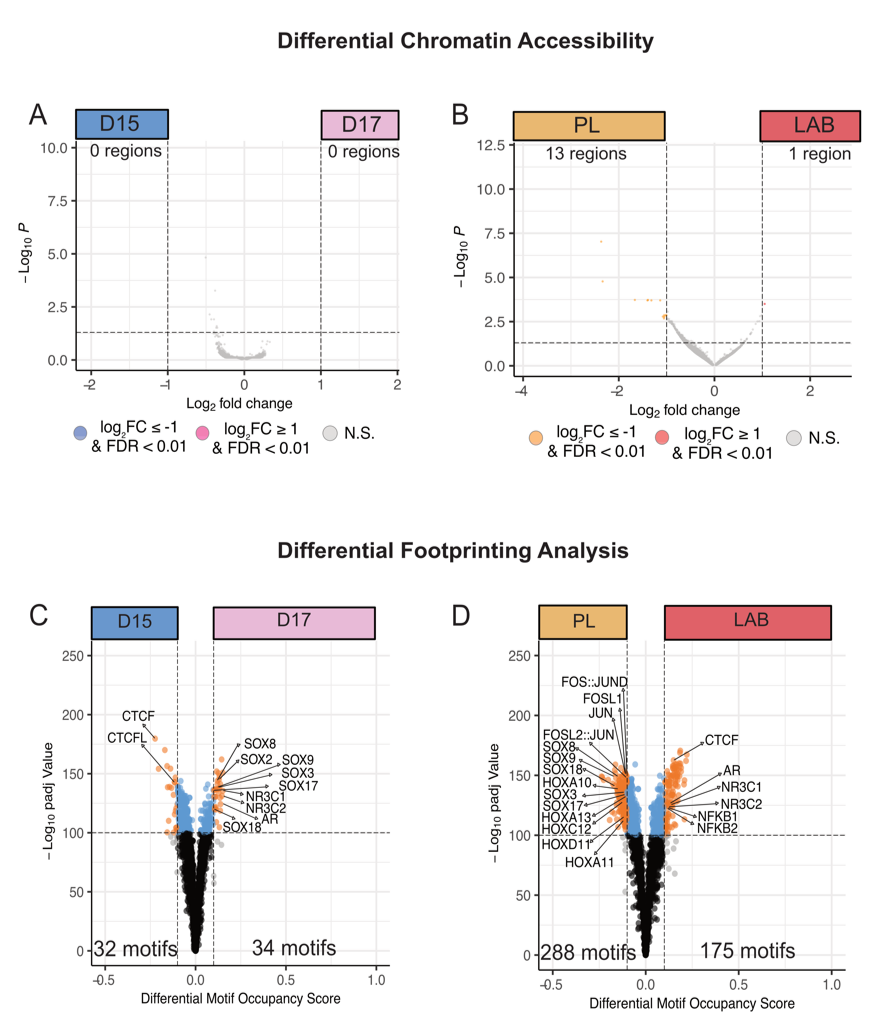

**Supplemental Figure 2: Pairwise comparisons of differential chromatin accessibility patterns between different timepoints of mouse gestation.**

Pairwise comparison in chromatin accessibility between A) day 15 (D15) and day 17 (D17); B) prelabor (day 19; PL) and labor (day 19.5; LAB). For all comparisons, |log_2_FC| ≥ 1 and FDR < 0.01 was used. Pairwise comparison in motif accessibility between C) day 15 (D15) and day 17 (D17); D) prelabor (day 19; PL) and labor (day 19.5; LAB).

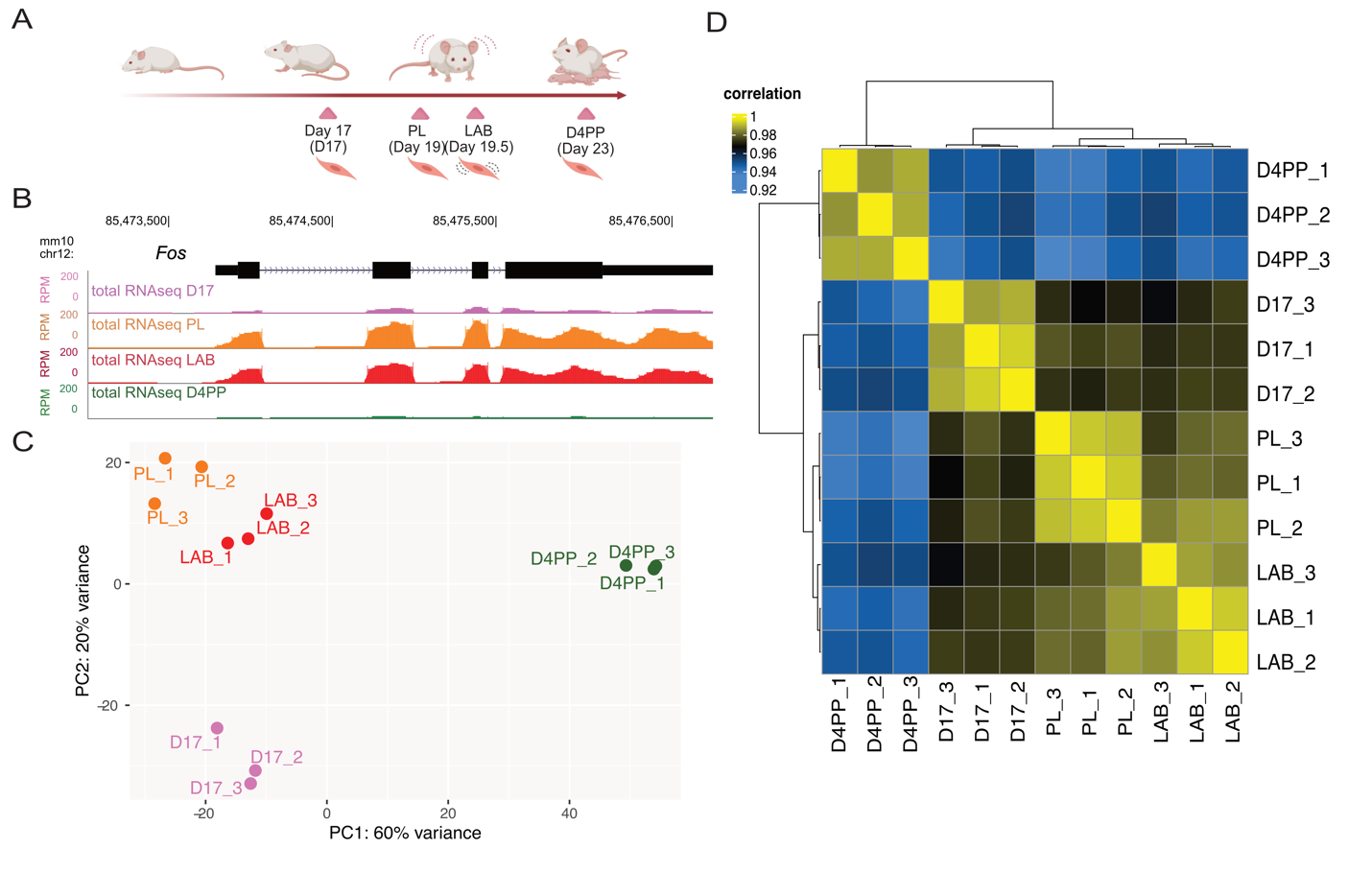
**Supplemental Figure 3: Gestational stage-specific RNA-seq replicates cluster based on timepoint of sample collections.**

A) Schematic of the mouse gestational timepoints outlining days when mouse myometrium tissue samples were collected for RNA-seq which include day 17 (D17) of late gestation, prelabor (PL; D19), active labor (LAB, D19.5), and day 4 postpartum (D23; D4PP) (n=3 per gestational timepoint). B) Total RNA-seq reads (reads per million; RPM) at the labor associated *Fos* gene locus demonstrating increased expression at the prelabor (PL) and labor (LAB) stages compared to late gestation (D17) and postpartum (D4PP). C) Principal component analysis (PCA) demonstrates clustering of RNA-seq replicates belonging to the same gestational timepoint. D) Hierarchical clustering of RNA-seq samples, where the lighter color indicates increased correlation, from D17, PL, LAB, and D4PP samples.

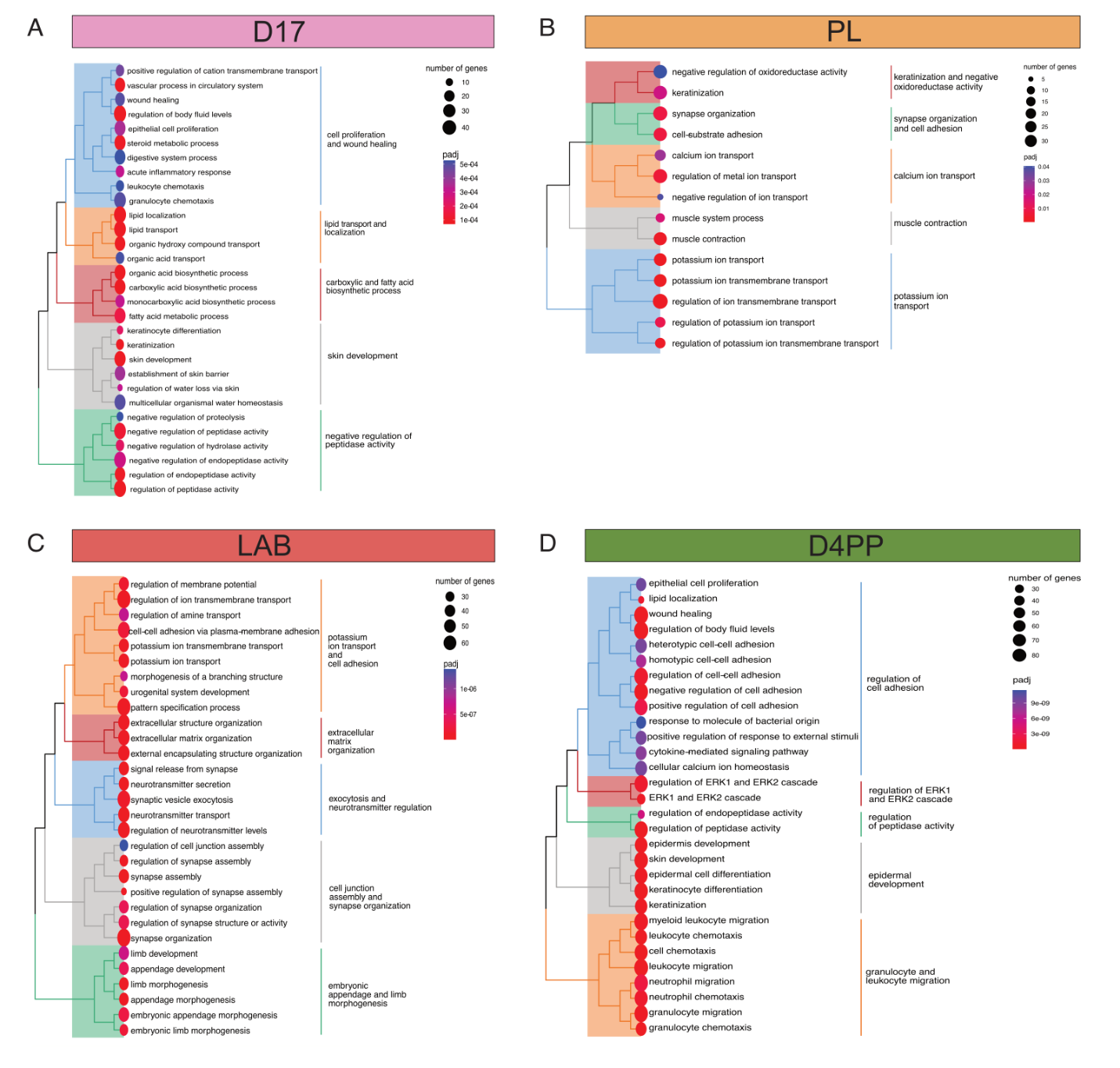

**Supplemental Figure 4: Genes expressed at different gestational stages are involved in distinct biological processes.**

A) Biological processes of genes expressed during pregnancy (D17), B) prelabor (PL), C) labor (LAB), or D) postpartum (D4PP).

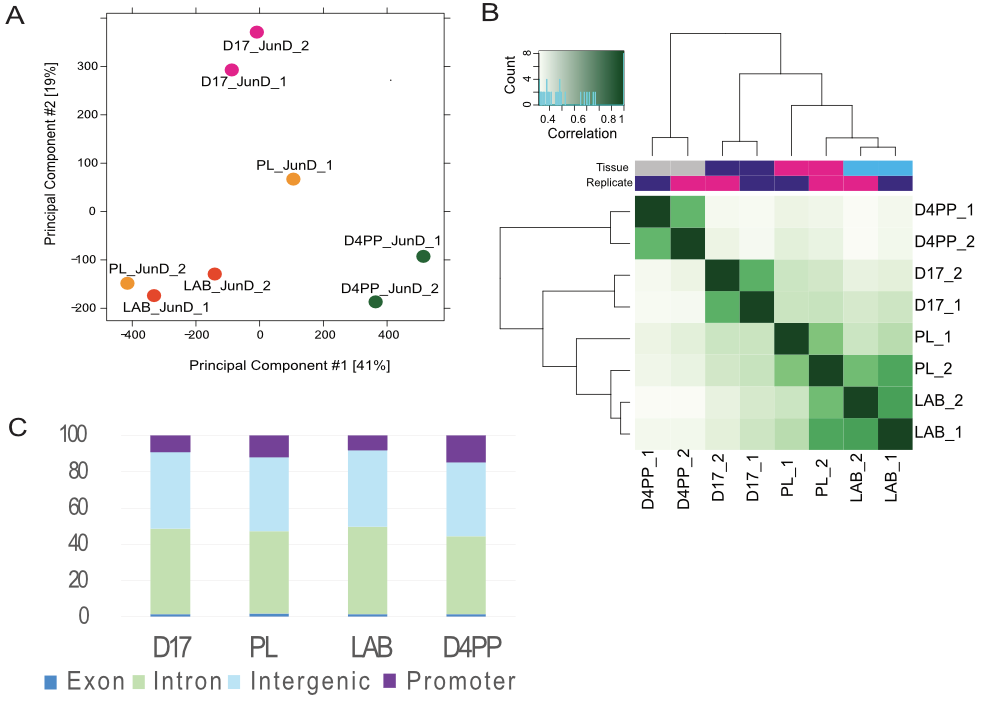

**Supplemental Figure 5: JUND ChIP-seq in mouse myometrium.**

A) Principal component analysis (PCA) demonstrating clustering of ChIP-seq replicates belonging to the same gestational timepoint. B) Hierarchical clustering of ChIP-seq samples, where the darker color indicates increased correlation, from D17, PL, LAB, and D4PP samples. C) Functional genomic annotation of JUND ChIP-seq peaks at D17, PL, LAB, and D4PP.

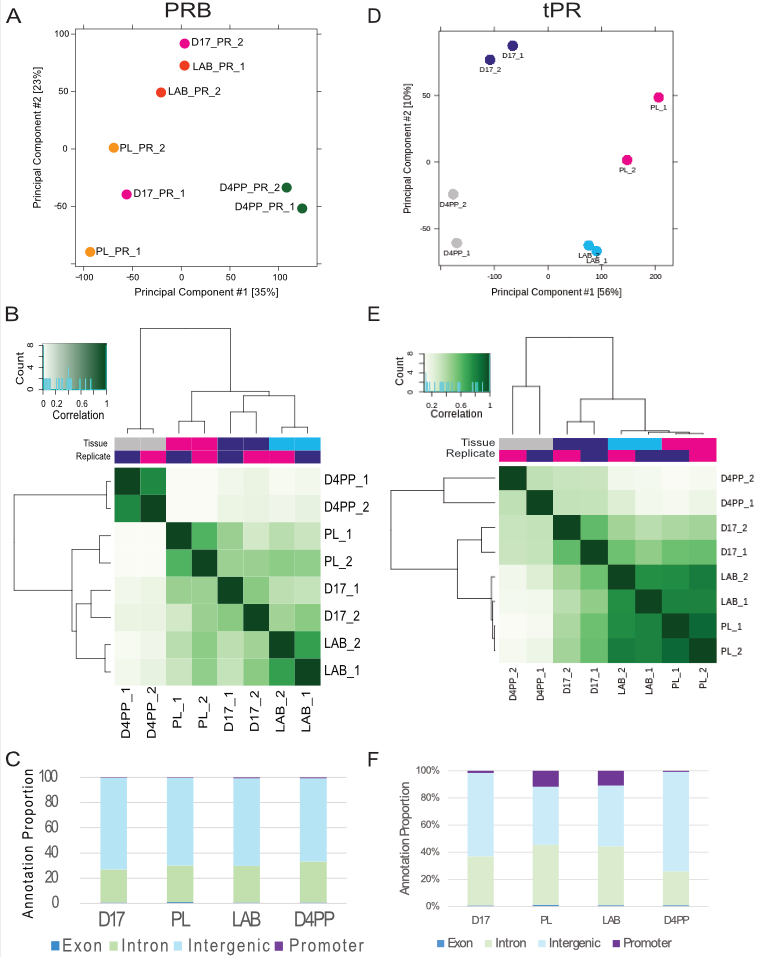

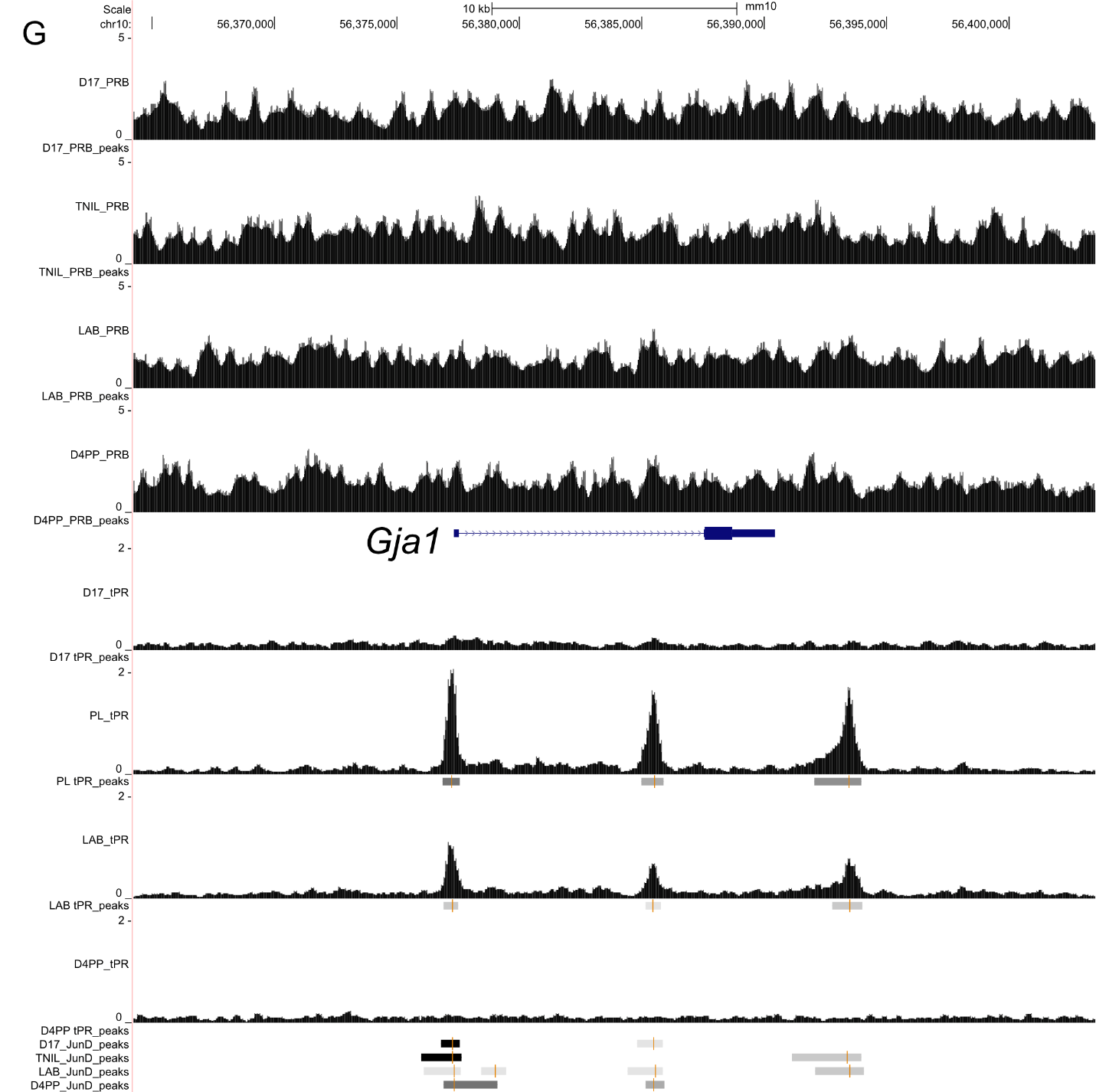

**Supplemental Figure 6: Total progesterone receptor (tPR) and progesterone receptor B (PRB) ChIP-seq in mouse myometrium.**  A) Principal component analysis (PCA) demonstrating clustering of PRB ChIP-seq replicates belonging to the same gestational timepoint. B) Hierarchical clustering of PRB ChIP-seq samples, where the darker color indicates increased correlation, from D17, PL, LAB, and D4PP samples. C) Functional genomic annotation of PRB ChIP-seq peaks at D17, PL, LAB, and D4PP. D) Principal component analysis (PCA) demonstrating clustering of tPR ChIP-seq replicates belonging to the same gestational timepoint. E) Hierarchical clustering of tPR ChIP-seq samples, where the darker color indicates increased correlation, from D17, PL, LAB, and D4PP samples. F) Functional genomic annotation of tPR ChIP-seq peaks at D17, PL, LAB, and D4PP. G) PRB and total PR (tPR) ChIP-seq enrichment at Gja1, no PRB peaks were detected (top), in the region with 3 strong tPR peaks (bottom).

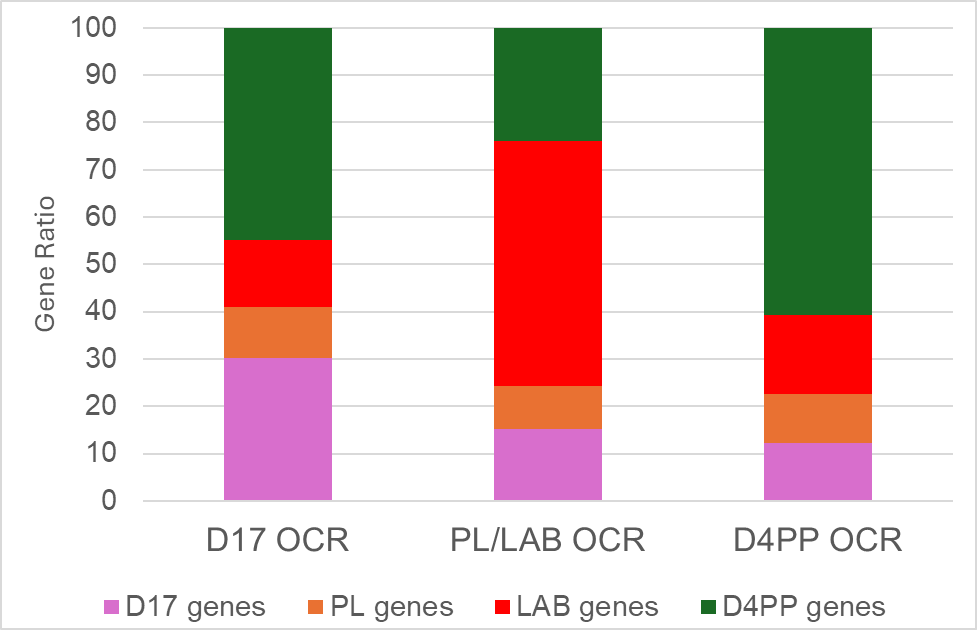

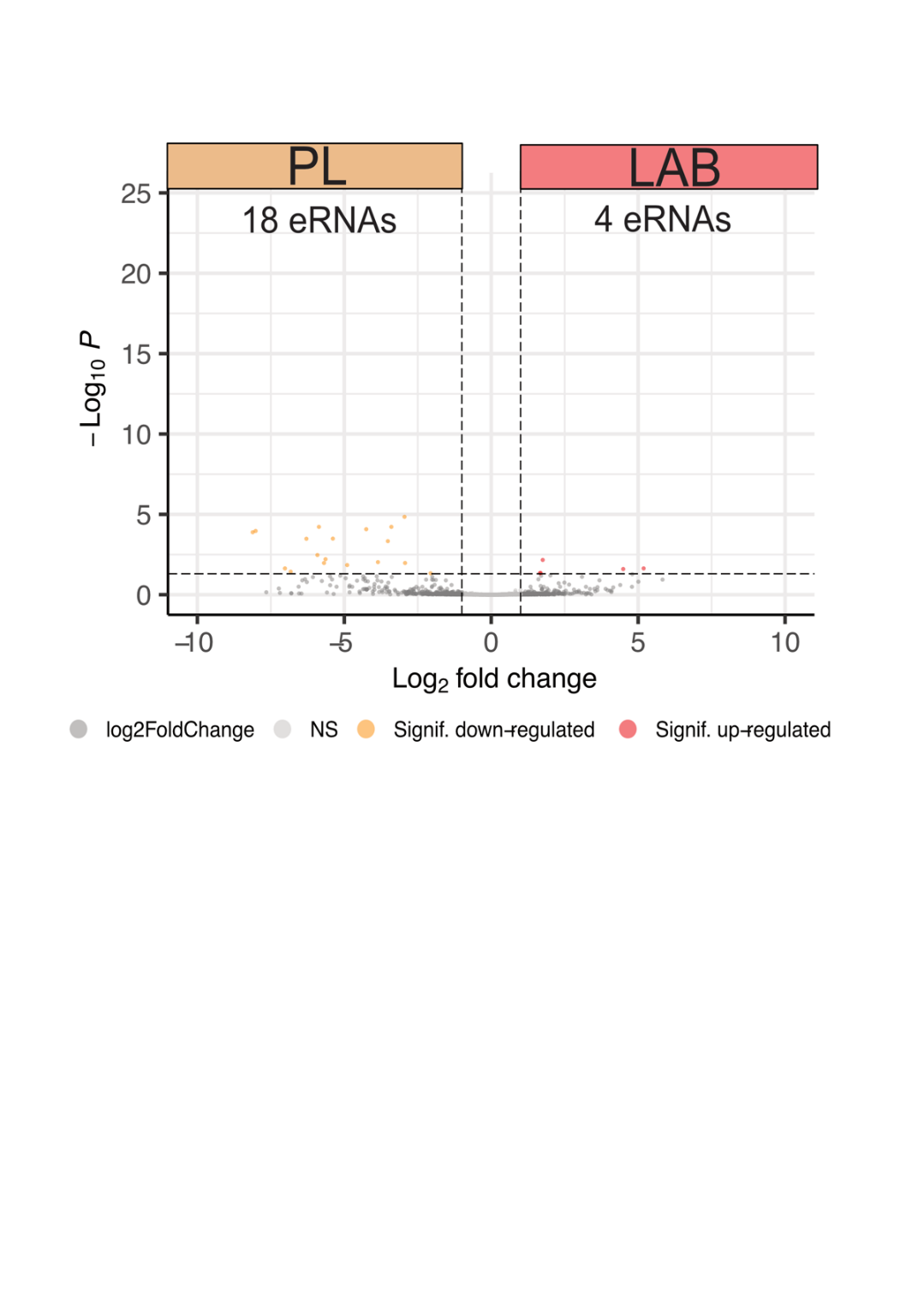

**Supplemental Figure 7: Gene expression groups associated with ATAC-seq open chromatin region (OCR) clusters and eRNAs dynamic between prelabor and labor.**

TOP: Stacked bar graph depicting the distribution of timepoint-specific genes that are associated with ATAC-seq (intronic or intergenic) peaks in each cluster. Genes demonstrating timepoint-specific expression common at both the intron and the exon level were used (from Fig 2F).  BOTTOM: Enhancer RNA (eRNA) expression at PL and LAB. Volcano plot displaying differential eRNA expression between between PL (prelabor) and labor (LAB).

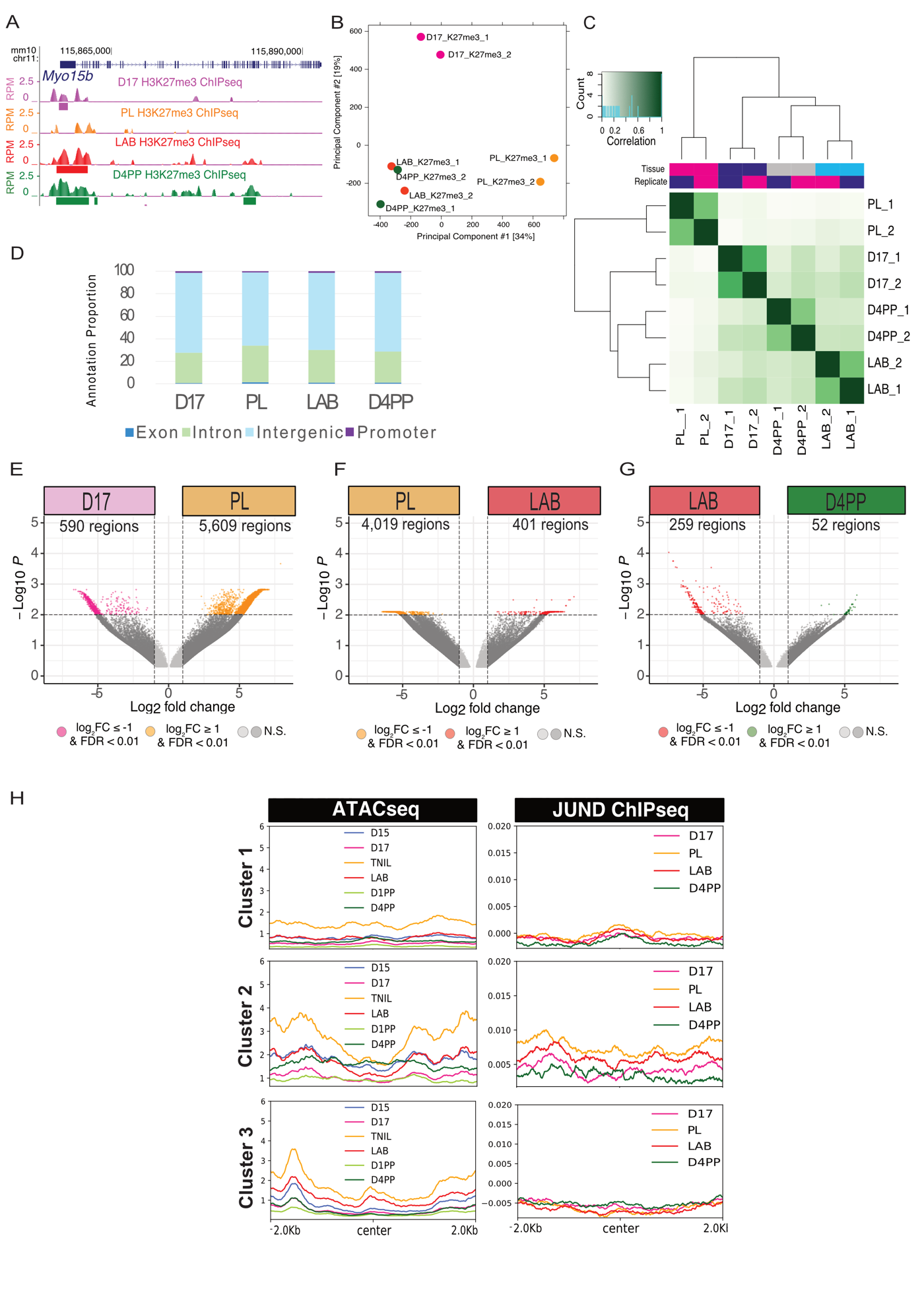

**Supplemental Figure 8: Increased H3K27me3 enrichment in the prelaboring (PL) myometrium.**

A) H3K27me3 ChIP-seq reads (reads per million; RPM) at the PL-upregulated *Myo15b* gene locus demonstrates high enrichment at all timepoints except prelabor (PL). B) Principal component analysis (PCA) demonstrates clustering of ChIP-seq replicates belonging to the same gestational timepoint. C) Hierarchical clustering of ChIP-seq samples, where the darker color indicates increased correlation, from D17, PL, LAB, and D4PP samples. D) Functional genomic annotation of H3K27me3 ChIP-seq peaks at D17, PL, LAB, and D4PP. Volcano plot analysis of pairwise comparisons depicting differential H3K27me3 enrichment between D17 and PL (E); PL and LAB (F); and LAB and D4PP (G). For all comparisons, |log_2_FC| ≥ 1, FDR < 0.01 was used. H) ATAC-seq and JUND ChIP-seq signal at the different H3K27me3 clusters.
